# Sortilin deficiency alters baseline retinal homeostasis and injury-induced signaling without affecting optic nerve crush-induced neurodegeneration

**DOI:** 10.64898/2026.05.08.723723

**Authors:** Thomas Stax Jakobsen, Anna Bøgh Lindholm, Toke Bek, Anders Nykjaer, Thomas J. Corydon, Anne Louise Askou

## Abstract

The effect of sortilin inhibition on acute inner retinal neurodegeneration induced by optic nerve crush was investigated. Pharmacological sortilin inhibition using intravitreal delivery of a polyclonal antibody or a small-molecule inhibitor was evaluated in C57BL/6JRj male mice subjected to unilateral crush. Inner retinal thickness was evaluated by optical coherence tomography, and retinal ganglion cell density was determined in retinal flat mounts. Furthermore, the effect of constitutive sortilin deficiency was examined using *Sort1-/-* mice. Changes in protein and mRNA levels of sortilin, p75^NTR^, and associated injury markers were analyzed.

Neither pharmacological inhibition or constitutive loss of sortilin protected against inner retinal thinning or retinal ganglion cell loss following optic nerve crush. A transient 1.4-fold increase in p75^NTR^ mRNA was observed early after injury, accompanied by a two-fold increase in protein levels. While sortilin expression remained largely unchanged, sortilin deficiency was associated with an altered baseline retinal state, including increased GFAP, p75^NTR^, and proBDNF levels. Following optic nerve crush, the induction of p75^NTR^ was significantly attenuated in sortilin-deficient retinas compared with wild type, without affecting the extent of RGC degeneration.

In summary, sortilin inhibition does not preserve inner retinal structure following optic nerve crush, but modulates glial activation, inflammatory signaling, and proneurotrophin dynamics. These findings indicate that sortilin-dependent pathways are not key drivers of optic nerve crush-induced neurodegeneration but may be more relevant in disease contexts characterized by chronic stress and neuroinflammation.

## 1 Introduction

The loss of retinal neurons, neurodegeneration, is a common feature of most visually debilitating retinal pathologies either as a primary mechanism or secondary to inflammatory and vascular insults. However, despite extensive efforts, translation of promising pre-clinical strategies into the clinic is currently lacking, and an unmet need exists for deeper understanding of involved mechanisms to identify novel targets.

The receptor sortilin (SORT1) is a member of the Vps10p domain family and has received interest as a therapeutic target in cardiovascular and metabolic disease^[1,2]^, cancer^[3,4]^, and dementias^[5]^. A neuroprotective effect of sortilin inhibition has mainly been linked to its role as a regulator of neurotrophin signaling^[6]^: Sortilin binds to the p75 neurotrophin receptor (p75^NTR^; also known as nerve growth factor receptor [NGFR]) forming a receptor complex necessary for induction of apoptosis and inflammation by pro-neurotrophins. However, modulation of the pleiotropic sorting functions of sortilin may also be important for neuroprotection. An example is the sortilin-mediated internalization and degradation of progranulin. Indeed, restoration of progranulin levels using sortilin inhibition is currently explored as a treatment for frontotemporal dementia caused by progranulin haploinsufficiency^[7]^.

Pro-neurotrophin engagement of p75^NTR^ has been amply implicated in outer and inner retinal neurodegeneration^[8,9]^. However, p75^NTR^ does not evidently localize to retinal ganglion cells (RGCs) themselves^[10]^, and the effect of the receptor on inner retinal neurodegeneration in the setting of ischemic retinopathies is dependent on paracrine mechanisms primarily involving Müller cells^[11,12]^. The effect of pro-neurotrophin stimulated p75^NTR^ signaling has been investigated in specific RGC loss models such as traumatic optic neuropathy, *i.e.* damage to the RGC axons in the optic nerve by crush or transection, or by induced ocular hypertension (OHT) mimicking glaucoma. Following optic nerve crush (ONC), increased p75^NTR^ protein levels and co-localization with the macroglia marker glial fibrillary acidic protein (GFAP) are observed^[13]^. Pharmacological inhibition of p75^NTR^ in Sprague–Dawley rats and p75^NTR^ knockout in mice have been reported to reduce RGC death following optic nerve transection (ONT)^[14]^. Similarly, small-molecule inhibition of p75^NTR^ protected against ONT and OHT in Wistar rats^[15]^, and optic nerve regeneration was also slightly increased in p75^NTR^ knockout mice following crush^[16]^. In contrast, others have observed no protective effect of pharmacological p75^NTR^ antagonism on RGC loss following OHT^[17]^.

The role of sortilin in retinal neuroprotection remains less well-characterized, but sortilin deficient mice show reduced rates of apoptosis in the developing retina^[18]^, sortilin loss is protective against light-dependent photoreceptor death *in vitro* and *in vivo*^[19]^, and sortilin upregulation in the retina of patients with diabetic retinopathy and a remarkable neuroprotective effect on the inner retina in streptozotocin-induced diabetic mice have been shown in our recent study^[20]^. In the central nervous system, sortilin-deficiency protects corticospinal neurons following injury^[18]^, but does not protect against neuronal cell death secondary to traumatic brain injury^[21]^. Regarding peripheral nerves, sortilin is dispensable for dorsal root ganglion cells loss following sciatic nerve injury^[22]^, and motor recovery is similar in knockout and wild-type (WT) mice although sortilin was suggested to modulate Schwann cell signaling and Remak bundle regeneration. On the other hand, sortilin contributes to the age-dependent degeneration of sympathetic neurons^[18]^. Thus, the ability to protect neurons seems highly cell and context dependent.

Given the established role of sortilin as a co-receptor for p75^NTR^ and regulator of proneurotrophin trafficking, inhibition of sortilin would be expected to attenuate p75^NTR^-dependent degenerative signaling following optic nerve injury. However, the contribution of sortilin to acute ONC-induced retinal degeneration remains unclear, particularly considering its pleiotropic roles in protein trafficking and glial biology. Moreover, it is not known whether sortilin influences key injury-associated pathways in the retina, including glial activation, inflammatory signaling, and proneurotrophin dynamics, in the context of ONC.

To address this, we investigated the effect of pharmacological sortilin inhibition and genetic sortilin deficiency on retinal structure, RGC survival, and injury-associated molecular responses following ONC.

## 2 Materials and methods

### 2.1 Animals

Male C57Bl/6JRj mice (Janvier, Le Genest-Saint-Isle, France) aged 8-10 weeks were used for examination of sortilin levels following ONC and therapeutic experiments with sortilin inhibitors. Male mice were chosen due to our previous implication of sortilin in diabetes-induced neurodegeneration in male mice^[20]^.

*Sort1-/-* animals were C57Bl/6J-Sort1^tm1Tew^/JBomTac mice originally created by replacing 126 bp from exon 14 and 303 bp of the subsequent intron sequence of sortilin with a neo-cassette using standard embryonic stem cell technology^[18]^. Homozygous *Sort1-/-* mice were used for breeding in-house and underwent regular backcrossing and genotyping. Age-matched C57Bl/6JBomTac mice (Taconic Biosciences, Germantown, New York) were used as WT controls.

Mice were kept on a 12h/12h light/dark cycle at the Animal Facilities at the Department of Biomedicine, Aarhus University, Denmark. Mice had ad libitum access to Altromin maintenance feed and water.

### 2.2 Anesthesia

Before surgery or non-invasive imaging, mice were anesthetized with an intraperitoneal (i.p.) injection of a mixture of ketamine (Ketador 60–100 mg/kg [Richter Pharma AG, Wels, Austria] and medetomidine hydrochloride (Cepetor 0.5–1 mg/kg [ScanVet Animal Health A/S, Fredensborg, Denmark]). Pupils were dilated with a drop of 1% tropicamide (Mydriacyl, Alcon Nordic A/S, Copenhagen, Denmark). During anesthesia, the eyes were lubricated with carbomer eye gel (Viscotears 2 mg/ml, Alcon Nordic). Immediately after procedures, anesthesia was reversed by atipamezole 0.5-1 mg/kg (Antisedan, Orion Pharma, Copenhagen, Denmark) and animals were placed on a heating plate until they moved spontaneously.

### 2.3 Optic nerve crush

ONC was performed in accordance with previous reports^[23,24]^ (Figure S1). Surgeries were performed in one eye under an OPMI 1 FR PRO Surgical microscope (Zeiss, Jena, Germany). A partial temporal peritomy was created with fine Vannas-Tübingen scissors (Fine Science Tools GmbH, Heidelberg, Germany) and the optic nerve was exposed by blunt dissection using Dumont angled forceps (Dumont, Montignez, Switzerland). The exposed optic nerve was crushed 1-2 mm behind the globe using Dumont #N7 curved, self-closing forceps for 5 seconds. Fundoscopy was performed using a coverslip to ensure patency of the central retinal artery. In the opposite eye, a sham-procedure was performed limited to a conjunctival peritomy and blunt dissection.

In preliminary experiments, fluorescein-angiography using i.p. injection of sodium fluorescein (Fluorescein 0.05 mg/g [Paranova Danmark A/S, Herlev, Denmark]) and fundus fluorescence imaging with the MICRON® IV imaging system (Phoenix research laboratories, Pleasanton, CA, USA) was performed to further ensure that the ONC method did not affect retinal perfusion.

### 2.4 Therapeutic compounds and intravitreal injection

Pharmaceutical sortilin inhibition was evaluated in two experiments. In the first experiment, the effect of 2 μL (1 μg/μL) intravitreally injected goat anti-sortilin polyclonal antibody (AF2934, R&D Systems, Minneapolis, MN, USA) was compared with normal goat IgG polyclonal antibody (AB-108-C, R&D Systems). Both antibodies were diluted in PBS (Biowest, Nuaillé, France). In the second experiment, the small-molecule inhibitor AF38469 (MedChemExpress LLC, Monmouth Junction, NJ, USA) was compared with the buffer solution. A stock solution of 100 μg/μL in DMSO (VWR Chemicals, Radnor, PA, USA) was diluted 1:100 in PBS. 2 μL of this 1 μg/μL was injected and a 1% DMSO in PBS solution was used as control.

Intravitreal injections were performed immediately following surgery in the eye subjected to ONC. The other eye was used as a non-injected control, *i.e.* subjected to the sham-procedure, but not intravitreally injected. A 30G disposable needle (B. Braun, Melsungen, Germany) was used to puncture the sclera near the limbus, and a 33G blunt-ended needle of a Hamilton syringe (Hamilton company, Reno, NV, USA) was then inserted into the opening and used to inject 2 μL of either therapeutic compound or control solution. The surgeon was blinded to treatment allocation during the procedure.

### 2.5 Optical coherence tomography

For evaluation of inner retinal neurodegeneration, circular peripapillary optical coherence tomography (OCT) B-scans were acquired using the MICRON® IV image-guided OCT 2 system (Phoenix research laboratories). Examinations were performed during the first two weeks following crush, where robust thinning of retinal layers occurs^[25]^. The average thickness of the NGI (retinal nerve fiber layer, ganglion cell, and inner plexiform layer) complex was determined by loading the OCT images into InSight software (Phoenix Research laboratories) and manually segmenting retinal layers. For characterization of retinal layers in *Sort1-/-* mice and WT additional horizontal, linear B-scans through the optic disc were acquired and segmented.

### 2.6 Flat mounting and immunostaining

Mice were sacrificed two weeks following crush surgery as robust loss of RGCs is present at this time point^[25]^. Following sacrifice, eyes were enucleated and fixed in 4% paraformaldehyde (VWR Chemicals) over-night at 4°C. The anterior segment and lens were removed and the neuroretina was carefully peeled off the retinal pigment epithelium (RPE)/choroid and immersed in PBS buffer. Before immunostaining, flat mounts were blocked and permeabilized over-night at 4°C in retina blocking buffer (RBB) containing PBS with 1% BSA (Sigma-Aldrich, St. Louis Burlington, MA, USA) and 0.5% Triton X100 (Sigma-Aldrich). Subsequently, the retinas were incubated with rabbit anti-RNA-binding protein with multiple splicing (RbPMS) 1-1.6 mg/ml (NBP2-20112, Novus Biologicals, LLC, Centennial, CO, USA) diluted 1:400-1:500 in RBB for 48 hours at 4°C. Next, the retinas were washed and incubated with Alexa-568 Donkey anti-rabbit 2 mg/ml (A10042, Thermo Fisher Scientific, Waltham, MA, USA) diluted 1:400 in RBB for 24 hours at 4°C. Following washing steps, the flat mounts were transferred to Super-Frost Plus glass slides (Thermo Fisher Scientific) and mounted with the RGC layer facing upwards using Fluoromount-G™ Mounting Medium (Thermo Fisher Scientific).

### 2.7 Wide-field fluorescence imaging and quantification of retinal ganglion cells

Image acquisition was performed at the Bioimaging Core Facility, Health, Aarhus University, Denmark. Retinal sections were imaged with the Olympus VS120 upright widefield fluorescence microscope (Olympus, Tokyo, Japan) equipped with Spectra X 7IR LED multi-spectral light engine and Semrock pentafilter (DAPI/FITC/Cy3/Cy5/Cy7 Penta LED HC Filter Set #F68-050) with

Hamamatsu ORCA-FLASH4.0 V2 (QE 82%) camera (Hamamatsu, Shizuoka, Japan). Images were taken with the Olympus UPlanSApo 20x/0.75 air objective and associated VS-ASW imaging software. Images were captured with fixed settings and processed similarly.

Quantification of RbPMS positive cells was performed in the QuPath 0.5.1 image analysis software^[26]^ using the automatic cell detection tool applied to whole retinas or by manual counting within a defined midperipheral area. Quantification was performed similarly within experimental cohorts.

### 2.8 Quantification of *Sort1* and *Ngfr* expression using RT-qPCR

Groups of C57Bl/6JRj mice were sacrificed without any intervention, or 3, 7, and 14 days post crush (dpc). Six mice were included in each group. In the control group sacrificed without intervention, one eye was used for Western blotting and one eye for qPCR. In the mice subjected to crush the contralateral eye served as sham. Eyes were dissected immediately and the neuroretina flash frozen and stored at -80°C until processing. For RNA extraction, neuroretinas were lysed using the RLT+ buffer (QIAGEN, Hilden, Germany) with 10 mM TCEP (Macherey-Nagel, Düren, Germany) and subsequently homogenized using a QIAshredder (QIAGEN). RNA was purified with the RNeasy micro kit according to protocol (QIAGEN). DNase treatment was performed using DNA-free™ Kit (Thermo Fisher Scientific) according to protocol (“routine DNase treatment”). The iScript cDNA synthesis kit (Bio-Rad, Hercules, CA, USA) was used for first-strand synthesis with 144 ng RNA as input.

qPCR reactions were prepared using RealQ Plus Master Mix Green (Amplicon, Odense, Denmark) using 1 µL cDNA diluted 1:20 and with a total reaction volume of 10 µL. Final primer concentrations were 1 µM. Reactions were run on the LightCycler 480 (Roche Diagnostics, Basel, Switzerland). A standard curve was prepared using a mix of cDNA samples in three-fold serial dilutions. Relative concentrations were calculated using the standard curve method using the LightCycler 480 software (Roche Diagnostics).

Primers used are presented in Supplementary Table 1. Glyceraldehyde 3-phosphate dehydrogenase (*Gapdh*) was used as endogenous control as it had been shown stable between ONC and sham groups in pilot experiments. Determined efficiencies were 109.5%, 93.1%, and 98.6% for *Sort1*, *Ngfr*, and *Gapdh*, respectively. Specificity was confirmed using melt curve analysis and/or gel electrophoresis. No-template and minus reverse transcriptase control Cq values were above the detection limit of Cq 35.

### 2.9 Quantification protein levels using Western blot

In an experiment evaluating changes in sortilin and p75^NTR^ following ONC, C57Bl/6JRj mice were sacrificed without any intervention, or 3, 7, and 14 dpc. Six mice were included in each group. In the control group sacrificed without intervention, one eye was used for Western blotting and one eye for qPCR. In the mice subjected to crush the contralateral eye served as sham. In an experiment evaluating effects of sortilin deficiency on protein changes following ONC, *Sort1-/-* and aged-matched C57Bl/6JBomTac mice were sacrificed without any intervention or 7 dpc. Five *Sort1-/-* and seven WT animals were used as baseline controls. Six mice of both genotypes were subjected to crush in one eye, while the contralateral eye served as sham.

Following sacrifice, eyes were dissected immediately and the neuroretina flash frozen and stored at -80°C until processing. For Western blot analysis, the neuroretinas were first thawed on ice. Subsequently, 100 µL radioimmunoprecipitation assay buffer (Thermo Fisher) with cOmplete™ Mini protease inhibitor cocktail (Roche Diagnostics) was added to each sample tube and tissue were homogenized using a metal bead and a Bullet Blender^®^ (Next Advance, Inc., Troy, NY, USA); samples were placed in the Bullet Blender for 30 seconds at 6000 rpm, then incubated 2 minutes on ice. This procedure was repeated twice if necessary. The homogenates were centrifuged at 13,000 *g* at 6°C for 10 minutes, after which the supernatant was transferred to a new microcentrifuge tube.

Protein concentrations were determined using the Bradford Assay Dye Reaction Concentrate (Bio-Rad) according to the manufacturer’s instructions, and 19-20 µg of total protein was loaded onto a 12% (sortilin, p75^NTR^, and anti-tumor necrosis factor α (TNFα)) or 4-15% (brain-derived neurotrophic factor (BDNF), GFAP, and p75^NTR^) Criterion^TM^ TGX Stain-Free^TM^ gel (Bio-Rad). Gels were run for 1 hour at 100 V and subsequently UV-activated to enable total protein visualization. Proteins were transferred onto a polyvinylidene difluoride membrane (Bio-Rad) using the Trans-Blot Turbo Transfer system (Bio-Rad) and total protein was visualized using the ChemiDoc Imaging system (Bio-Rad). Membranes were blocked for 1 hour at room temperature in Tris-buffered saline (Thermo Fisher Scientific) with 0.1% Tween20 (Sigma-Aldrich) (TBS-T) containing 5% w/v skim-milk powder (VWR Chemicals). Membranes were incubated at 4°C over-night with anti-sortilin antibodies (ab16640, 1.0 mg/ml, Abcam, Cambridge, UK) 1:1000, anti-nerve growth factor receptor (NGFR) antibodies (AF1157, 0.2 mg/ml, R&D Systems) 1:1000, anti-TNFα antibodies (ab6671, 1 mg/ml, Abcam) 1:2000, anti-BDNF antibodies (abcam108319, 0.28 mg/ml, Abcam) 1:5000, or anti-GFAP antibodies (ab5804, 2 mg/ml, Millipore, Merck Life Science A/S, Søborg, Denmark) 1:5000. For Figure 4, anti-NGFR antibodies (07-476, 1 mg/ml, Millipore) were used at a dilution of 1:1000 to detect p75^NTR^. After a 3x5 min wash in TBS-T, the membranes were subsequently incubated 1 hour at room temperature with HRP-conjugated secondary antibodies diluted 1:10,000 (Goat-anti-rabbit, Bio-Rad; Rabbit-anti-goat, 0.5 g/l DAKO, Agilent Technologies, Santa Clara, CA). Clarity^TM^ Western ECL Substrate (179-5060, Bio-Rad) was used for detection, and imaging was performed with the ChemiDoc MP imaging system (Bio-Rad). Quantification was performed using Image Lab Software (Bio-Rad). To account for potential variability between gels and enable accurate comparison of protein expression, control samples were included on all gels as inter-gel controls. Signal intensities from these common samples were used to normalize data across gels.

**Figure 1.**
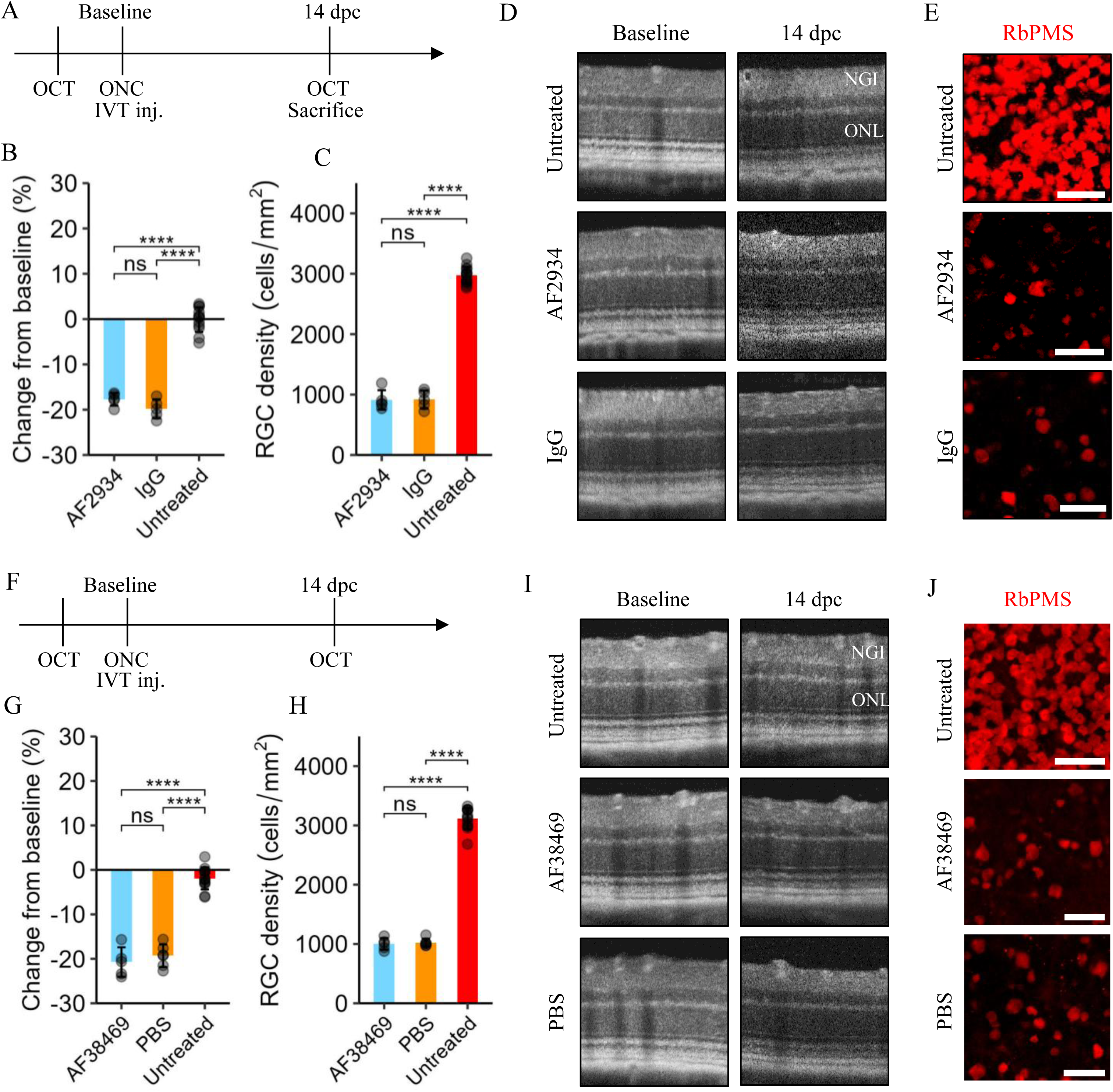
Sortilin inhibition using a polyclonal antibody or small-molecule inhibitor does not preserve inner retinal structure. A. Experimental plan: Following baseline optical coherence tomography (OCT) scans, C57BL6/JRj mice were subjected to optic nerve crush (ONC) in one eye and sham procedure in the other eye. Immediately after, mice were randomized to receive intravitreal (IVT) injection of either the anti-sortilin polyclonal antibody (pAb) AF2934 or IgG control (2 µL of 1 µg/µL) in the crush eye while the other served as untreated sham control. Mice were sacrificed following OCT at 14 days post crush (dpc). B. Bar plot (mean ± sd) showing quantification of the change from baseline of the combined thickness of the nerve fiber, ganglion cell and inner nuclear layer, denoted the NGI thickness. Statistical comparisons were performed using one-way ANOVA followed by Tukey’s test (n > 5). C. Bar plot (mean ± sd) showing quantification of the density of RNA-binding protein with multiple splicing (RbPMS) positive retinal ganglion cells (RGCs) in retinal flat mounts. Statistical comparisons were performed by one-way ANOVA followed by Tukey’s test (n > 5). Representative OCT scans (D) and areas from anti-RbPMS (red) immunostained retinal flat mounts (E) are shown. Scale bars 50 µm. F. Experimental plan: As in (A), C57BL6/JRj mice were randomized to receive IVT injection of the small-molecule inhibitor AF38469 (2 µL of 1 µg/µL) or vehicle (PBS-1% DMSO) in the crush eye. G. Bar plot (mean ± sd) showing quantification of the change from baseline of NGI thickness. Statistical comparisons were performed by one-way ANOVA followed by Tukey’s test (n > 5). H. Bar plot (mean ± sd) showing quantification of RGC density in retinal flat mounts. Statistical comparisons were performed by one-way ANOVA followed by Tukey’s test (n > 5). Representative OCT scans (I) and areas from anti-RbPMS (red) immunostained retinal flat mounts (J) are shown. Scale bars 50 µm. Significance levels: * [0.01, 0.05]; ** [0.001,0.01]; *** [0.0001, 0.001]; **** [0, 0.0001].

**Figure 2.**
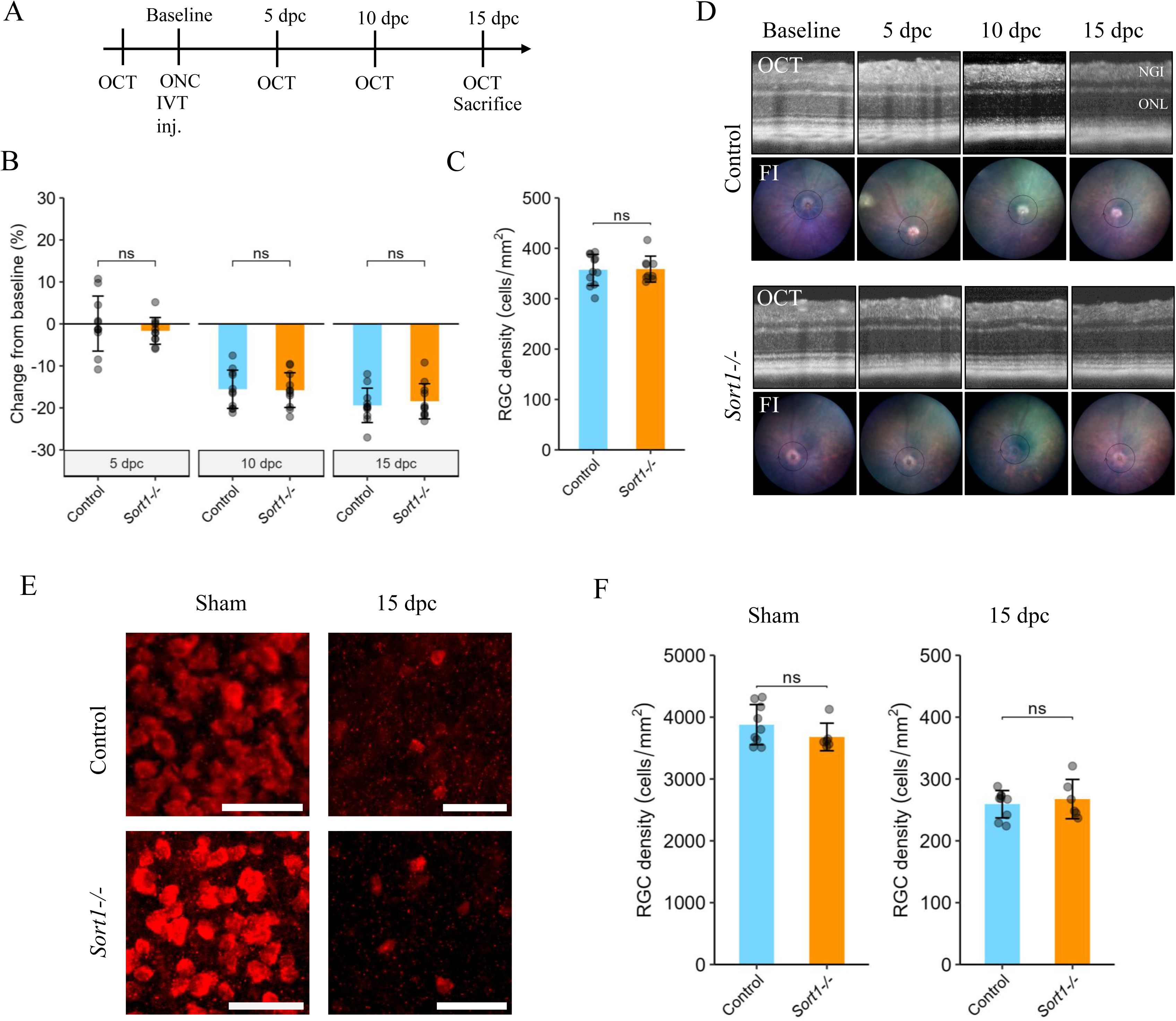
Sortilin deficiency does not affect inner retinal pathology following optic nerve crush. A. Experimental plan: 9-week-old *Sort1-/-* mice and age-matched C57Bl/6JBomTac wild-type mice were subjected to optic nerve crush (ONC) in the right eye. Optical coherence tomography (OCT) scans were performed at baseline and 5, 10, and 15 days post crush (dpc). B. Bar plot (mean ± sd) showing quantification of change from baseline of the combined thickness of the nerve fiber, ganglion cell, and inner nuclear layer denoted the NGI thickness. Statistical comparisons were performed using linear mixed-effects model to account for repeated measurements (n = 10-11). C. Bar plot (mean ± sd) showing quantification of RNA-binding protein with multiple splicing (RbPMS) positive retinal ganglion cells (RGCs) 15 dpc. Statistical comparison was performed using Student’s t-test (n = 10-11). D. Representative fundus images (FI) and OCT scans at the respective dpc. E. Representative areas from anti-RbPMS (red) immunostained retinal flat mounts in a separate replication cohort. Scale bars 50 µm. F. Bar plots (mean ± sd) showing quantification of RbPMS positive RGCs 15 dpc in eyes subjected to crush (right) and the contralateral sham eyes (left). Statistical comparisons were performed using Student’s t-test (n = 6-9). Significance levels: * [0.01, 0.05]; ** [0.001,0.01]; *** [0.0001, 0.001]; **** [0, 0.0001].

**Figure 3.**
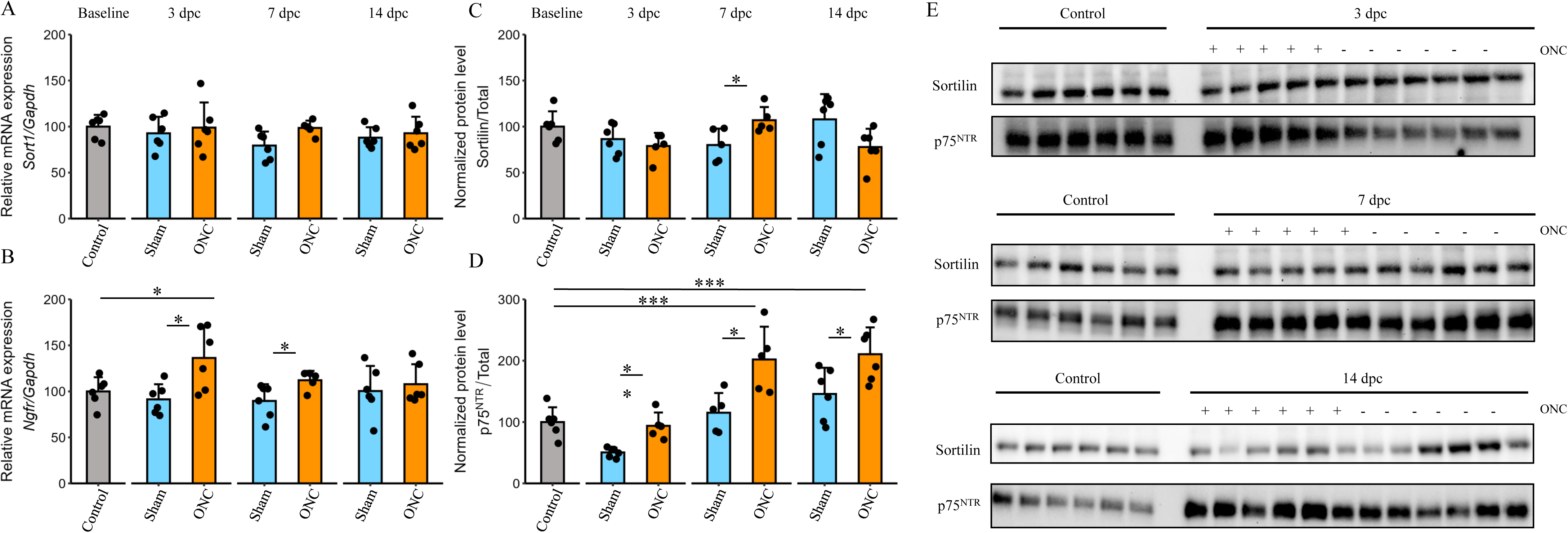
Sortilin and p75^NTR^ expression following optic nerve crush. Neuroretinal samples from control eyes not subjected to any procedure as well as eyes subjected to optic nerve crush (ONC) or sham procedure at 3, 7, and 14 days post crush (dpc) were analyzed A. and B. Bar plots showing expression of *Sort1* and *Ngfr* related to the endogenous control *Gapdh*. C and D. Bar plots (mean ± sd) showing quantification of sortilin and p75^NTR^ protein levels related to total protein. E. Western blots used for quantification. Statistical comparisons of the difference between control and ONC group at different time points and comparisons between ONC and sham group at different time points were performed using two- or one-way ANOVA followed by Tukey’s test and Dunnet’s test respectively (n = 5-6 at each time point). Significance levels: * [0.01, 0.05]; ** [0.001,0.01]; *** [0.0001, 0.001]; **** [0, 0.0001].

**Figure 4.**
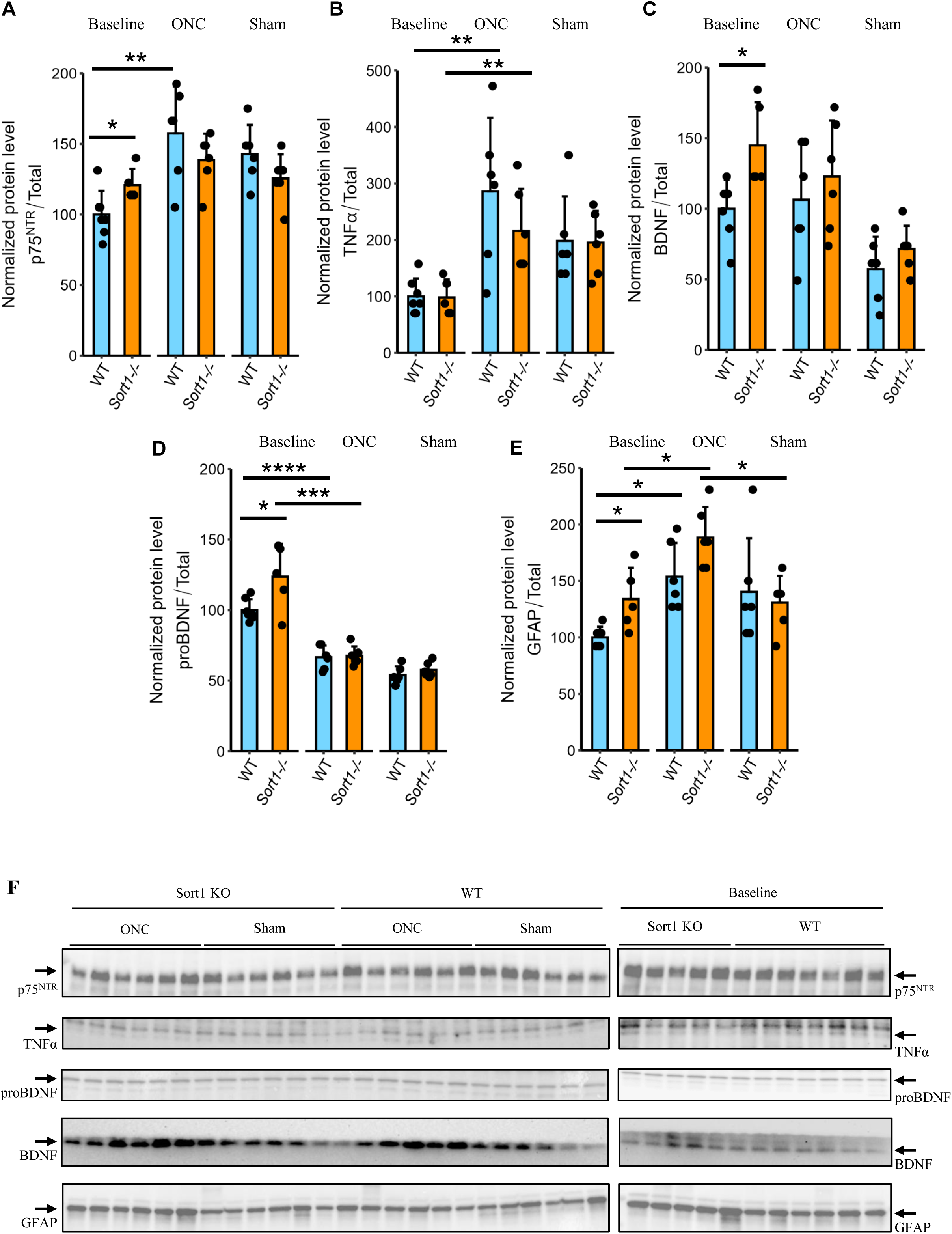
Influence of sortilin deficiency on protein levels following optic nerve crush. Neuroretinal samples from *Sort1-/-* and age-matched C57Bl/6JBomTac wild-type (WT) mice not subjected to any procedure or with eyes subjected to optic nerve crush (ONC) or sham procedure and sacrificed 7 days post crush (dpc) were analyzed. A-E. Bar plots (mean ± sd) showing quantification of protein levels related to total protein in Western blots for p75^NTR^, TNFα, proBDNF, BDNF, and GFAP, respectively. F. Western blots used for quantification. Statistical comparisons of the difference between WT and *Sort1-/-* animals in the different groups and the difference from baseline levels were performed using two-way ANOVA followed by Tukey’s test (n = 5-7). Significance levels: * [0.01, 0.05]; ** [0.001,0.01]; *** [0.0001, 0.001]; **** [0, 0.0001].

### 2.10 Statistical analysis

Values are presented as mean ± sd unless stated otherwise. Statistical analyses were performed using the R statistical software package R-4.4.1 (https://www.r-project.org/). A significance level of α = 0.05 was used and confidence intervals are presented accordingly.

Data was assessed for normality by QQ-plot inspection. The exact analyses are specified in figure legends. Linear mixed-effects models for repeated measurements were implemented using the *lmer* package.

## 3 Results

### 3.1 Sortilin inhibition using a polyclonal antibody fails to preserve inner retinal thickness or retinal ganglion cell density following optic nerve crush

The polyclonal AF2934 antibody block nerve growth factor (NGF) and sortilin interaction as shown by the ability to inhibit pro-NGF stimulated neurite elongation^[27]^. We have earlier observed a protective effect of intravitreally injected AF2934 on inner retinal neurodegeneration in streptozotocin-induced diabetic mice^[20]^. Furthermore, this antibody has been shown to protect mice from mechanically induced allodynia^[28]^. We thus wanted to investigate whether intravitreal injection of this antibody could protect against inner retinal neurodegeneration induced by ONC.

Following baseline OCT measurements, ONC was performed in one eye and sham surgery in the other eye. This was followed immediately by intravitreal injection in the eye subjected to ONC. OCT measurements were performed 14 dpc, immediately before sacrifice and harvest of eyes for flat mounting and quantification of RGCs (Figure 1A). A total of 15 8-week-old animals were randomized to receive either the therapeutic AF2934 antibody or IgG control (2 µL of 1 µg/µL). Reduction in NGI thickness from baseline measurements was not significantly different being 2.1% [-1.69; 5.91] (p = 0.36) smaller in the AF2934 group compared with the control group. Likewise, RGC density in flat mounts was not significantly different with -6 cells/mm^2^ [-239;228] (p = 0.998) in the AF2934 group compared with the control group (Figure 1B-E).

To investigate whether a higher dose would confer benefit, a separate experiment was performed using a higher dose (2 µL of 5 µg/µL) and with quantification of NGI thickness in OCT scans acquired 5, 10, and 15 dpc. 12 mice were included in each group. Mice were randomized to receive either AF2934 or IgG control. A greater NGI thickness reduction from baseline amounting to -2.8% [-5.2;0.3] (p = 0.034) was observed in the high-dose AF2934 group compared with the IgG group at 3 dpc, but otherwise no difference was observed (Figure S2).

### 3.2 Sortilin inhibition using the small-molecule inhibitor AF38469 does not preserve inner retinal thickness or retinal ganglion cell density following optic nerve crush

The small-molecule inhibitor AF38469^[29]^ blocking the interaction between sortilin and the NGF pro-domain^[30]^ has been used to study *e.g.* the cancer promoting effects of sortilin^[3]^. Therefore, the therapeutic potential of intravitreal AF38469 delivery in the ONC model was evaluated: Following baseline OCT measurements, ONC was performed in one eye and sham surgery in the other eye. This was followed immediately by intravitreal injection in the eye subjected to ONC. OCT measurements were performed 14 dpc, immediately before sacrifice and harvest of eyes for flat mounting and quantification of RGCs (Figure 1F). A total of 15 8-week-old animals were randomized to receive either AF38469 or vehicle control (2 µL of 1 µg/µL). No therapeutic effect was observed: Reduction in NGI thickness from baseline measurements was not significantly different being 1.41% [- 2.66;5.49] (p = 0.66) larger in the AF38469 group compared with the control group. Likewise, RGC density in flat mounts was not significantly different with -22 cells/mm^2^ [-243;200] (p = 0.996) in the AF38469 group compared with the control group (Figure 1G-J).

### 3.3 Sortilin deficiency does not protect the inner retina following optic nerve crush

Pharmacological inhibition may be highly dependent on dose and timing. Thus, to confirm our findings, we also performed experiments in *Sort1-/-* mice. Sortilin-deficient mice are viable and without known adverse phenotypes. Sortilin-deficient mice show reduced neuronal loss during development.^[18]^ However, they have normal fundoscopic appearance and quantification of retinal layers in OCT scans acquired from sortilin-deficient and age-matched WT animals did not reveal structural differences (Figure S3). These findings indicate preserved retinal structure in adulthood, although potential adaptive changes during development cannot be excluded, as described for *Ngfr- /-* mice^[31]^.

Following baseline OCT measurements ONC was performed in the right eye and sham surgery in the left eye in cohorts of sortilin-deficient and age-matched WT animals. Twelve animals were included in each group; two and one animal, respectively, were sacrificed prematurely due to humane endpoints. Follow-up OCT was performed 5, 10, and 15 dpc. Following sacrifice 15 dpc, eyes were harvested for flat mounting and quantification of RGC density (Figure 2A,D). Change in NGI thickness from baseline was not significantly different at any time point between sortilin-deficient and WT control mice being 1.0% [-2.7;4.7] (p = 0.59) smaller in the sortilin-deficient mice (Figure 2B). Similarly, RGC densities in retinal flat mounts were similar between the two groups. The difference between the sortilin-deficient and the WT control mice was 2 cells/mm^2^ [-25;28] (p = 0.996) (Figure 2C). The RGC density 15 days following ONC was considerably lower than in experiments using C57Bl/6JRj mice. Hence, the quantification of RGC density was replicated in a separate cohort including six sortilin-deficient animals and nine age-matched WT controls: Observed RGC densities were similar and with no difference between sortilin-deficient and WT control mice 15 dpc. In addition, RGC densities in the sham eyes were similar between the groups (Figure 2E,F) and comparable to those in C57Bl/6JRj.

### 3.4 Regulation of sortilin and p75^NTR^ following optic nerve crush

Changes in ocular sortilin expression have not previously been evaluated following ONC, but an ∼1.5 fold increase in p75^NTR^ mRNA in the first days following ONC in mice^[16]^ as well as an increase in p75^NTR^ protein have been reported^[13]^. To identify neuroretinal alterations in sortilin and p75^NTR^ mRNA and protein following ONC, groups of male C57Bl/6JRj mice were sacrificed before any intervention (8-weeks-old) and 3, 7, and 14 dpc. Successful crush was confirmed by uniform NGI thickness reduction on OCT scans (Figure S4).

Relative neuroretinal *Sort1* mRNA expression (Figure 3A) was not significantly different in eyes subjected to ONC compared with controls. Similarly, no significant differences were observed compared with baseline levels. *Ngfr* expression was significantly increased 3 and 7 dpc compared with controls and the contralateral sham-treated eye but decreased to normal levels 14 dpc (Figure 3B). The increase at 3 dpc amounted to 44.9% [8.9;81.0] (p = 0.021).

Western blotting showed a small significant 27.0% (p = 0.03) increase in sortilin protein levels in the ONC eyes compared with controls 7 dpc (Figure 3E), but no increase relative to baseline at any time point (Figure 3C). The p75^NTR^ protein level was significantly increased in retinas from eyes subjected to ONC 7 and 14 dpc compared with baseline and the contralateral sham eye (Figure 3D). The increase relative to baseline amounted to 101.98% [43.1;160.9] (p < 0.0001) at 7 dpc and 110.6% [54.4;166.8] (p = 0.0003) at 14 dpc. Complete blots are provided as supplemental information (Figure S5).

### 3.5 Influence of sortilin deficiency on protein levels following optic nerve crush

Western blot analysis was performed to assess protein levels of p75^NTR^, TNFα, BDNF, and GFAP in whole retinal lysates from WT and sortilin-deficient mice under baseline conditions, and 7 days following ONC or sham surgery (Figure 4).

At baseline, sortilin-deficient retinas exhibited significantly increased levels of p75^NTR^ (20.8% [1.47;40.0] (p = 0.037)) and GFAP (33.8% [6.71;61.0] p = 0.02) compared to WT, indicating differences in basal protein expression. ProBDNF (∼32 kDa) and BDNF (∼15 kDa) levels were also elevated in sortilin-deficient retinas (23.7% [3.09;44.3], p = 0.0283 and 44.9% [12.2;77.6], p = 0.012, respectively), whereas TNFα levels were comparable between genotypes (−2% [-43.2;39.2], p = 0.916).

Following ONC, p75^NTR^ levels increased significantly in WT retinas (57.5% [26.2;88.8], p=0.00193), while the small increase in sortilin-deficient mice was insignificant (17.8% [-3.96;39.5], p = 0.0973) The increase was significantly larger in WT animals (39.1% [16.4;61.7], p = 0.017). A similar pattern was observed for TNFα and GFAP, where protein levels increased after ONC in both WT (186% [74.5;297], p = 0.00366) and sortilin-deficient (118% [36.1;200], p = 0.0098) retinas. A tendency toward a less pronounced increase in sortilin-deficient retinas was observed although not significantly (−65.6 [-203;71.8], p = 0.312).

ProBDNF levels decreased following ONC in both WT (−33.4 [-43.2;-23.7], p < 0.0001) and sortilin-deficient (−56.2 % [-78.4;-33.9], p = 0.0003) retinas to similar levels (0.97% [-8.76;10.7], p = 0.829) The decrease was slightly larger in sortilin-deficient retinas (−12.0 [-21.0;-2.95], p = 0.0144). No consistent genotype-dependent differences were observed in mature BDNF levels. Complete blots are provided as supplemental information (Figure S6).

## 4. Discussion

In the present study we did not observe a protective effect of pharmacological sortilininhibition or constitutive sortilin deficiency on inner retinal thinning or RGC loss following ONC. However, sortilin deficiency was associated with a distinct alteration of the retinal baseline state and partial modulation of injury response. At baseline, sortilin-deficient retinas exhibited increased levels of GFAP, p75^NTR^, and proBDNF indicating altered glial activation and proneurotrophin signaling. Following ONC, both WT and sortilin-deficient retinas showed robust injury responses, but the increase in p75^NTR,^ was attenuated in sortilin-deficient retinas. Similarly, a tendency of an attenuated TNFα response was observed. Despite these molecular differences, RGC loss and structural degeneration were comparable between genotypes.

Evaluation of neuroretinal expression of neurotrophins and their receptors in optic nerve injury models are somewhat limited, and this is the first investigation of changes in sortilin levels following ONC. The lack of robust alterations at both the transcript and protein level are compatible with lack of structural rescue although changes in cellular localization or cell-specific expression patterns cannot be excluded from the available data. Our finding of a transient increase in transcript levels of *Ngfr* is comparable with the ∼1.5 fold increase observed the first days following ONC in a mouse study.^[16]^ Similarly, the increase in p75^NTR^ protein observed in retinas from eyes subjected to ONC is in line with a study showing an ∼3-fold increase in p75^NTR^ protein most pronounced 14 days following ONC in Long Evans rats^[13]^. The difference in magnitude may depend on the specific experimental procedure, animal species, and protein quantification including normalization to total protein in our study instead of a single housekeeping protein. Based on established models of proneurotrophin signaling, in which proBDNF engages the p75^NTR^/sortilin receptor complex to promote apoptotic and inflammatory responses, increased proBDNF and p75^NTR^ levels would be expected to enhance degenerative signaling^[32]^. However, despite elevated baseline levels of proBDNF and p75^NTR^, sortilin deficiency did not result in increased RGC loss, suggesting that increased ligand and receptor availability alone is insufficient to drive degenerative signaling in the absence of sortilin. Conversely, the injury-induced increase in TNFα appeared attenuated in sortilin-deficient retinas without improving survival. Together, these observations suggest that modulation of the proneurotrophin/p75^NTR^ axis is not limiting for ONC-induced degeneration in this model.

Sortilin is an enigmatic receptor with pleiotropic functions^[33]^. A major focus for the therapeutic use of sortilin inhibition is the dependence on sortilin for pro-neurotrophin signaling through p75^NTR^ ^[6]^. Activation of the sortilin/p75^NTR^ receptor complex can induce apoptosis^[6]^ or inflammation^[10]^ depending on the cell type. In the eye, p75^NTR^ is not evidently present on adult RGCs^[10]^. Rather the protective effect of blocking p75^NTR^ in models of acute and chronic neurodegeneration seems to rely on inhibiting p75^NTR^ activation on Müller cells. Hence, intraocular injection of pro-NGF induces RGC loss with upregulation of TNFα expression in Müller cells, and this effect is blocked by sortilin deficiency^[11]^. In models of acute and chronic optic nerve injury, p75^NTR^ signaling promotes inner retinal neurodegeneration^[34]^: p75^NTR^ antagonism or knockout have been shown to ameliorate RGC loss following ONT^[14,15]^ and OHT^[15]^ as well as in induced diabetes^[12]^, and these effects are dependent on non-autonomous mechanism^[12,14]^.

We recently showed that inhibition of the p75^NTR^ co-receptor sortilin also confer neuroprotection in induced diabetes^[20]^ and sought in the present study to examine if this would translate into a protective effect in a model of acute inner retinal neurodegeneration. However, we did not observe any effect using pharmacological inhibition or knockout of sortilin under the experimental conditions applied. A true effect may exist outside the effect size that our experiments were powered to identify, and the pharmacological effect may be sensitive to the chosen agent and its pharmacokinetics^[14]^. However, replication using different approaches suggests that any existing effect is not robust. The absence of neuroprotection likely reflects the nature of the ONC model, in which RGC degeneration is driven by rapid, intrinsic injury-induced pathways that may proceed independently of glial-mediated and proneurotrophin-dependent signaling. In this context, sortilin-dependent pathways influence the retinal environment but are insufficient to alter the course of ONC-induced cell death. The loss of RGC in optic nerve injury and glaucoma has been hypothesized to depend on lack of neurotrophic support, which may decrease following disrupted axonal transport^[34]^. The neurotrophin BDNF and its cognate receptor TrkB seem to play a particularly important role^[35]^. Hence, it is interesting that sortilin also regulates BDNF secretion and degradation in a complex way^[36–38]^ and that it in certain contexts enhances Trk receptor transport to facilitate neurotrophin signaling^[39]^. In line with this, sortilin deficiency was associated with increased proBDNF levels without a corresponding increase in mature BDNF, indicating altered neurotrophin processing rather than enhanced neurotrophic support. This contrasts with studies demonstrating neuroprotection following exogenous BDNF or TrkB activation^[34,40–42]^ and suggests that endogenous changes in BDNF-related signaling in sortilin-deficient retinas are not sufficient to modify degeneration in this model.This study did not investigate functional measures such as electroretinography or optomotor responses, but these must be expected to correlate with the structural damage and be severely impaired in the used model system^[43]^. An effect on optic nerve regeneration was not specifically investigated, which could be pursued in further studies.

In summary, sortilin inhibition does not preserve inner retinal structure following ONC. However, sortilin influences glial activation, inflammatory signaling, and proneurotrophin dynamics in the retina. These effects do not translate into neuroprotection in the ONC model but may be more relevant in disease settings characterized by chronic stress and sustained neuroinflammation, such as diabetic retinopathy.

## Ethical statement

Animals were handled in accordance with the “Statement for the Use of Animals in Ophthalmic and Vision Research” from the Association for Research in Vision and Ophthalmology (ARVO). All animal experiments were performed under the approval of The Danish Animal Inspectorate (Case# 2020-15-0201-00556 [approval date 09-06-2020]).

## Author contributions

The authors confirm contribution to the paper as follows: Study conception and design: TS Jakobsen, AL Askou, and TJ Corydon; resources: TS Jakobsen, AL Askou, TJ Corydon, and AL Nykjaer; data collection: TS Jakobsen, AB Lindholm, and AL Askou; analysis and interpretation of results: TS Jakobsen, AB Lindholm, AL Askou, and TJ Corydon; draft manuscript preparation: TS Jakobsen; draft review and editing: TS Jakobsen, AB Lindholm, AL Askou, TJ Corydon; supervision: AL Askou, TJ Corydon, and T Bek. All authors reviewed the results and approved the final version of the manuscript.

## Acknowledgements

The authors thank Tina Hindkjær for expert technical assistance, and Stella Solveig Nolte for assistance with breeding of the knockout animals. The Aarhus University Health Bioimaging Core Facility is thanked for the use of equipment and technical assistance. The Animal Facility, Department of Biomedicine, Aarhus University, is thanked for housing the mice.

This work was supported by the Faculty of Health Sciences, Aarhus University (PhD scholarships to T.S.J. and A.B.L.), Fight for Sight, Denmark (T.S.J.), Ophthalmologist Else Bruntse’s foundation (T.S.J.), the Synoptik Foundation (T.S.J.), the A.P. Moller Foundation (T.S.J), the Dagmar Marshalls Foundation, the Independent Research Fund Denmark (grant no. 2034-00036B [T.J.C]), the Riisfort foundation (A.L.A), the Danish National Research Foundation (grant no. DNRF133 [A.N.]), and the Lundbeck Foundation (grant no. R480-2024-1094).

## Data availability

The datasets used and/or analyzed during the present study are available from the corresponding author upon reasonable request.

## References

1. Goettsch C, Kjolby M, Aikawa E. 2018. Sortilin and Its Multiple Roles in Cardiovascular and Metabolic Diseases. *Arteriosclerosis*, Thrombosis, and Vascular Biology 38:19–25

2. Møller PL, Rohde PD, Winther S, Breining P, Nissen L, et al. 2021. Sortilin as a Biomarker for Cardiovascular Disease Revisited. Front Cardiovasc Med 8:652584

3. Rhost S, Hughes É, Harrison H, Rafnsdottir S, Jacobsson H, et al. 2018. Sortilin inhibition limits secretion-induced progranulin-dependent breast cancer progression and cancer stem cell expansion. Breast Cancer Research 20

4. Yang W, Wu P-F, Ma J-X, Liao M-J, Wang X-H, et al. 2019. Sortilin promotes glioblastoma invasion and mesenchymal transition through GSK-3β/β-catenin/twist pathway. Cell Death & Disease 10

5. Salasova A, Monti G, Andersen OM, Nykjaer A. 2022. Finding memo: versatile interactions of the VPS10p-Domain receptors in Alzheimer’s disease. Molecular Neurodegeneration 17

6. Nykjaer A, Lee R, Teng KK, Jansen P, Madsen P, et al. 2004. Sortilin is essential for proNGF-induced neuronal cell death. Nature 427:843–48

7. Rhinn H, Tatton N, McCaughey S, Kurnellas M, Rosenthal A. 2022. Progranulin as a therapeutic target in neurodegenerative diseases. Trends in Pharmacological Sciences 43:641–52

8. Malik SC, Sozmen EG, Baeza-Raja B, Le Moan N, Akassoglou K, Schachtrup C. 2021. In vivo functions of p75NTR: challenges and opportunities for an emerging therapeutic target. Trends in Pharmacological Sciences 42:772–88

9. Mirzahosseini G, Adam JM, Nasoohi S, El-Remessy AB, Ishrat T. 2022. Lost in Translation: Neurotrophins Biology and Function in the Neurovascular Unit. Neuroscientist:10738584221104982

10. Garcia TB, Hollborn M, Bringmann A. 2017. Expression and signaling of NGF in the healthy and injured retina. Cytokine & Growth Factor Reviews 34:43–57

11. Lebrun-Julien F, Bertrand MJ, De Backer O, Stellwagen D, Morales CR, et al. 2010. ProNGF induces TNFα-dependent death of retinal ganglion cells through a p75NTRnon-cell-autonomous signaling pathway. Proceedings of the National Academy of Sciences 107:3817–22

12. Barcelona PF, Sitaras N, Galan A, Esquiva G, Jmaeff S, et al. 2016. p75NTR and Its Ligand ProNGF Activate Paracrine Mechanisms Etiological to the Vascular, Inflammatory, and Neurodegenerative Pathologies of Diabetic Retinopathy. The Journal of Neuroscience 36:8826–41

13. Mesentier-Louro L, De Nicolò S, Rosso P, De Vitis L, Castoldi V, et al. 2017. Time-Dependent Nerve Growth Factor Signaling Changes in the Rat Retina During Optic Nerve Crush-Induced Degeneration of Retinal Ganglion Cells. International Journal of Molecular Sciences 18:98

14. Lebrun-Julien F, Morquette B, Douillette A, Saragovi HU, Di Polo A. 2009. Inhibition of p75(NTR) in glia potentiates TrkA-mediated survival of injured retinal ganglion cells. Mol Cell Neurosci 40:410–20

15. Bai Y, Dergham P, Nedev H, Xu J, Galan A, et al. 2010. Chronic and acute models of retinal neurodegeneration TrkA activity are neuroprotective whereas p75NTR activity is neurotoxic through a paracrine mechanism. J Biol Chem 285:39392–400

16. Fujita Y, Takashima R, Endo S, Takai T, Yamashita T. 2011. The p75 receptor mediates axon growth inhibition through an association with PIR-B. Cell Death & Disease 2:e198–e98

17. Shi Z, Birman E, Saragovi HU. 2007. Neurotrophic rationale in glaucoma: A TrkA agonist, but not NGF or a p75 antagonist, protects retinal ganglion cellsin vivo. Developmental Neurobiology 67:884–94

18. Jansen P, Giehl K, Nyengaard JR, Teng K, Lioubinski O, et al. 2007. Roles for the pro-neurotrophin receptor sortilin in neuronal development, aging and brain injury. Nature Neuroscience 10:1449–57

19. Santos AM, López-Sánchez N, Martín-Oliva D, De La Villa P, Cuadros MA, Frade JM. 2012. Sortilin Participates in Light-dependent Photoreceptor Degeneration in Vivo. PLOS ONE 7:e36243

20. Jakobsen TS, Østergaard JA, Kjolby M, Birch EL, Bek T, et al. 2023. Sortilin Inhibition Protects Neurons From Degeneration in the Diabetic Retina. Invest Ophthalmol Vis Sci 64:8

21. Staib-Lasarzik I, Gölz C, Bobkiewiecz W, Somnuke P, Sebastiani A, et al. 2024. Sortilin is dispensable for secondary injury processes following traumatic brain injury in mice. Heliyon 10:e35198

22. Gürgör P, Pallesen LT, Johnsen L, Ulrichsen M, de Jong IE, Vaegter CB. 2016. Neuronal death in the dorsal root ganglion after sciatic nerve injury does not depend on sortilin. Neuroscience 319:1–8

23. Suter T, Wang J, Meng H, He Z. 2021. Utilizing mouse optic nerve crush to examine CNS remyelination. STAR Protoc 2:100796

24. Cameron E, Xia X, Galvao J, Ashouri M, Kapiloff M, Goldberg J. 2020. Optic Nerve Crush in Mice to Study Retinal Ganglion Cell Survival and Regeneration. BIO-PROTOCOL 10

25. Pang I-H, Clark AF. 2020. Inducible rodent models of glaucoma. Progress in Retinal and Eye Research 75:100799

26. Bankhead P, Loughrey MB, Fernández JA, Dombrowski Y, McArt DG, et al. 2017. QuPath: Open source software for digital pathology image analysis. Scientific Reports 7

27. Kalous A, Nangle MR, Anastasia A, Hempstead BL, Keast JR. 2012. Neurotrophic actions initiated by proNGF in adult sensory neurons may require peri-somatic glia to drive local cleavage to NGF. Journal of Neurochemistry 122:523–36

28. Richner M, Pallesen LT, Ulrichsen M, Poulsen ET, Holm TH, et al. 2019. Sortilin gates neurotensin and BDNF signaling to control peripheral neuropathic pain. Science Advances 5:eaav9946

29. Schrøder TJ, Christensen S, Lindberg S, Langgård M, David L, et al. 2014. The identification of AF38469: an orally bioavailable inhibitor of the VPS10P family sorting receptor Sortilin. Bioorg Med Chem Lett 24:177–80

30. Malik I, Christensen S, Stavenhagen JB, Dietz GPH. 2018. Development of a Cell-Based Assay to Assess Binding of the proNGF Prodomain to Sortilin. Cellular and Molecular Neurobiology 38:827–40

31. Harada C, Harada T, Nakamura K, Sakai Y, Tanaka K, Parada LF. 2006. Effect of p75NTR on the regulation of naturally occurring cell death and retinal ganglion cell number in the mouse eye. Developmental Biology 290:57–65

32. Teng HK, Teng KK, Lee R, Wright S, Tevar S, et al. 2005. ProBDNF induces neuronal apoptosis via activation of a receptor complex of p75NTR and sortilin. J Neurosci 25:5455–63

33. Nykjaer A, Willnow TE. 2012. Sortilin: a receptor to regulate neuronal viability and function. Trends in Neurosciences 35:261–70

34. Almasieh M, Wilson AM, Morquette B, Cueva Vargas JL, Di Polo A. 2012. The molecular basis of retinal ganglion cell death in glaucoma. Progress in Retinal and Eye Research 31:152–81

35. Mysona BA, Zhao J, Bollinger KE. 2017. Role of BDNF/TrkB pathway in the visual system: therapeutic implications for glaucoma. Expert Review of Ophthalmology 12:69–81

36. Evans SF, Irmady K, Ostrow K, Kim T, Nykjaer A, et al. 2011. Neuronal Brain-derived Neurotrophic Factor Is Synthesized in Excess, with Levels Regulated by Sortilin-mediated Trafficking and Lysosomal Degradation. Journal of Biological Chemistry 286:29556–67

37. Chen Z-Y, Ieraci A, Teng H, Dall H, Meng C-X, et al. 2005. Sortilin Controls Intracellular Sorting of Brain-Derived Neurotrophic Factor to the Regulated Secretory Pathway. The Journal of Neuroscience 25:6156–66

38. Yang M, Lim Y, Li X, Zhong J-H, Zhou X-F. 2011. Precursor of Brain-derived Neurotrophic Factor (proBDNF) Forms a Complex with Huntingtin-associated Protein-1 (HAP1) and Sortilin That Modulates proBDNF Trafficking, Degradation, and Processing. Journal of Biological Chemistry 286:16272–84

39. Vaegter CB, Jansen P, Fjorback AW, Glerup S, Skeldal S, et al. 2011. Sortilin associates with Trk receptors to enhance anterograde transport and neurotrophin signaling. Nature Neuroscience 14:54–61

40. Mansour-Robaey S, Clarke DB, Wang YC, Bray GM, Aguayo AJ. 1994. Effects of ocular injury and administration ofbrain-derived neurotrophic factor on survival and regrowth of axotomized retinalganglion cells. Proceedings of the National Academy of Sciences 91:1632–36

41. Di Polo A, Aigner LJ, Dunn RJ, Bray GM, Aguayo AJ. 1998. Prolonged delivery of brain-derived neurotrophic factor by adenovirus-infected Müller cells temporarily rescues injured retinal ganglion cells. Proceedings of the National Academy of Sciences 95:3978–83

42. Cheng L, Sapieha P, Kittlerova P, Hauswirth WW, Di Polo A. 2002. TrkB gene transfer protects retinal ganglion cells from axotomy-induced death in vivo. J Neurosci 22:3977–86

43. Liu Y, McDowell CM, Zhang Z, Tebow HE, Wordinger RJ, Clark AF. 2014. Monitoring retinal morphologic and functional changes in mice following optic nerve crush. Invest Ophthalmol Vis Sci 55:3766–74

