## Supplementary for "Sortilin deficiency alters baseline retinal homeostasis and injury-induced signaling without affecting optic nerve crush-induced neurodegeneration"

### Supplementary material

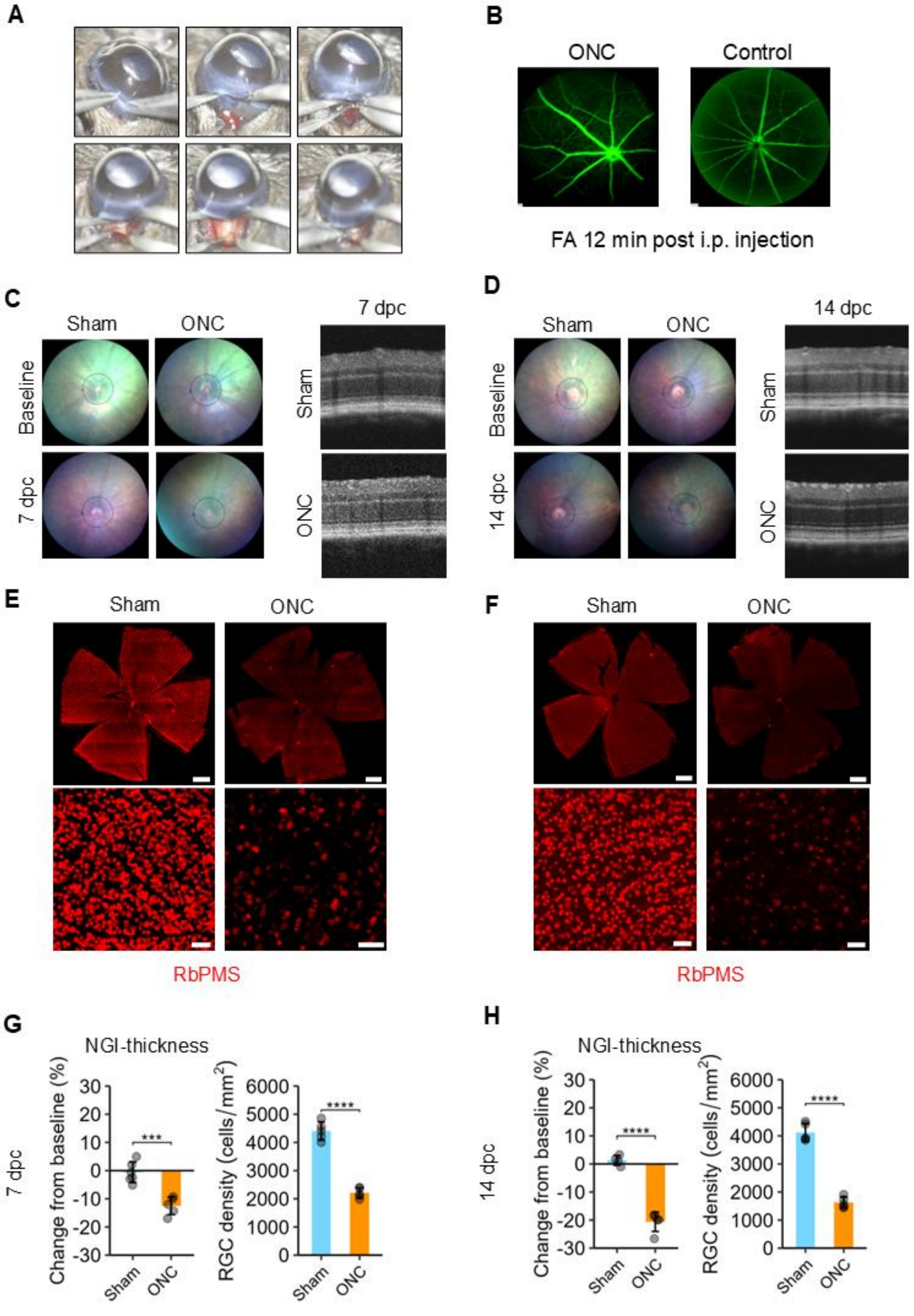
A
B
ONC
Control
FA 12 min post i.p. injection
C
Sham
ONC
Baseline
7 dpc
7 dpc
Sham
ONC
D
Sham
ONC
Baseline
14 dpc
14 dpc
Sham
ONC
E
Sham
ONC
RbPMS
F
Sham
ONC
RbPMS
G
NGI-thickness
7 dpc
Change from baseline (%)
RGC density (cells/mm²)
***
****
Sham
ONC
H
NGI-thickness
14 dpc
Change from baseline (%)
RGC density (cells/mm²)
****
****
Sham
ONC

**Figure S1. Validation of optic nerve crush model.** A. Surgical method of optic nerve crush (ONC): A peritomy is performed and the optic nerve is exposed by blunt dissection. The optic nerve is crushed approximately 1 mm behind the globe using self-closing forceps with pressure applied for 5 seconds. B. Fluorescein angiography (FA) demonstrating perfusion of retinal vessels following crush surgery. C. Representative fundus images and peripapillary circumferential optical coherence tomography (OCT) scans in C57BL/6JRj mice subjected to crush (left eye) or sham (right eye) and sacrificed 7 days post crush (dpc). D. Representative fundus images and OCT scans in mice sacrificed 14 dpc. E,F. Retinal flat mounts stained with antibodies against RNA-binding protein with multiple splicing (RbPMS) from the eye presented in C,D. Scalebars: 500 µm (overview) and 50 µm (magnification). G,H. Bar plots (mean ± sd) with quantification of the change in the combined thickness (NGI-thickness) of the nerve fiber layer, the ganglion cell layer, and the inner plexiform layer from baseline as well as the density of RbPMS positive ganglion cells in retinal flat mounts; statistical comparisons are performed with unpaired t-tests. Significance levels: * [0.01, 0.05]; ** [0.001,0.01]; *** [0.0001, 0.001]; **** [0, 0.0001].

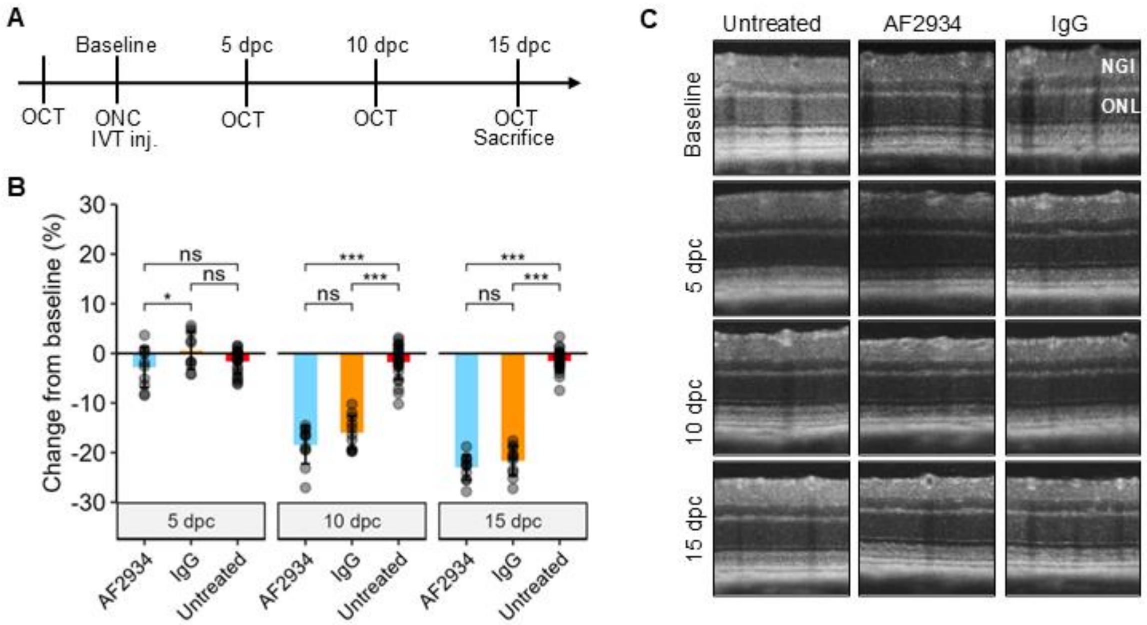

**Figure S2. Sortilin inhibition using a higher dose of polyclonal antibody does not preserve inner retinal thickness.** A. Experimental plan: Following baseline optical coherence tomography (OCT) scans, C57BL6/JRj mice were subjected to optic nerve crush (ONC) in one eye and sham procedure in the other eye. Immediately after mice were randomized to receive intravitreal (IVT) injection of either the anti-sortilin polyclonal antibody AF2934 or IgG control (2 µL of 5 µg/µL) in the crush eye while the other served as untreated sham control. Mice were sacrificed followed by OCT until sacrifice 15 days post crush (dpc). B. Bar plot (mean ± sd) showing quantification of NGI thickness. Statistical comparisons were performed using a linear mixed-effects model with eye nested under animal as the included random effect (n > 10 at each timepoint). C. Representative OCT scans at baseline and respective dpc. Significance levels: * [0.01, 0.05]; ** [0.001,0.01]; *** [0.0001, 0.001]; **** [0, 0.0001].

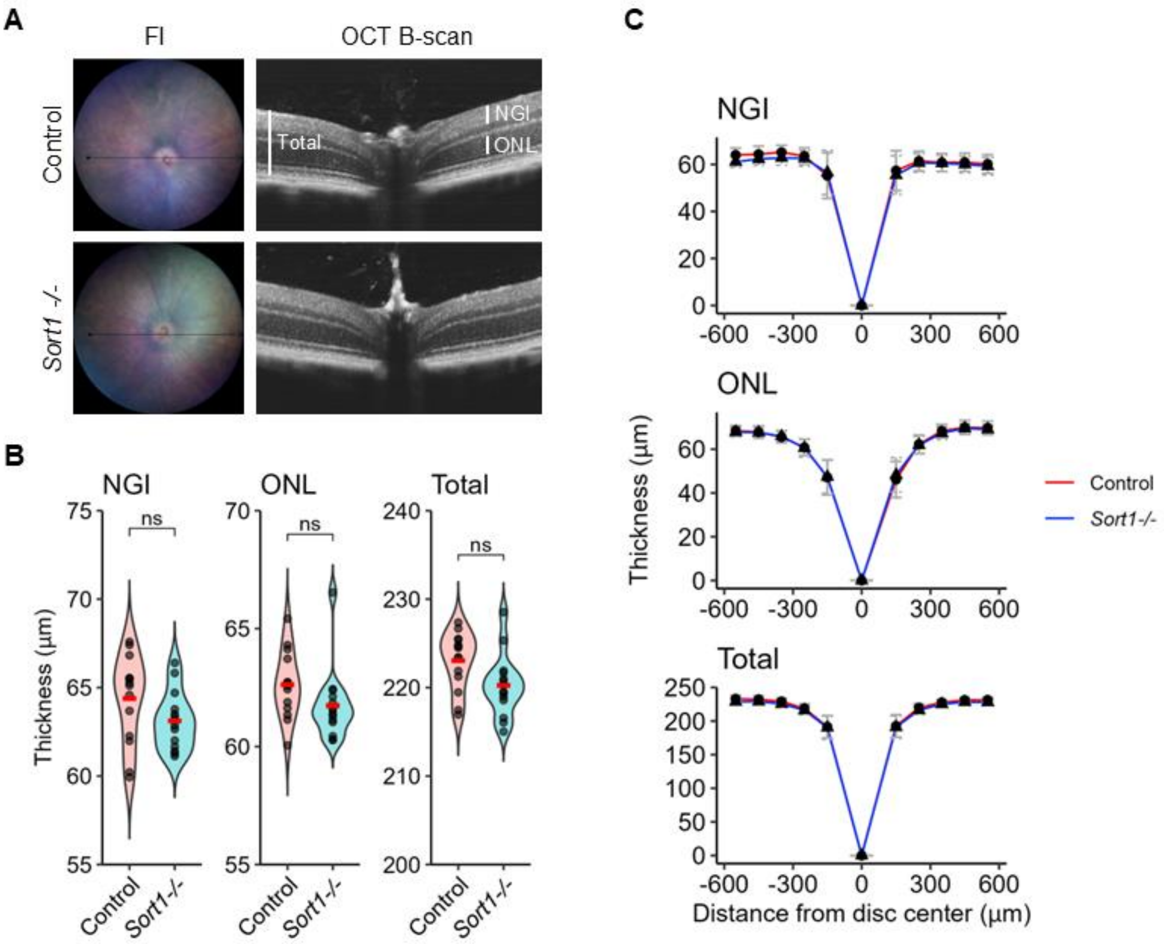

**Figure S3. Retinal structure of *Sort1-/-* (C57Bl/6J-Sort1tm1Tew/JBomTac) mice and age-matched C57Bl/6JBomTac wild-type mouse.** A. Representative fundus images (FI) and optical coherence tomography (OCT) B-scans. White bars mark the NGI (nerve fiber, ganglion cell, and inner plexiform layer), the outer nuclear layer (ONL) and total retinal thickness. B. Violin plot of NGI, ONL, and total retinal thickness in control and *Sort1-/-* animals quantified in peripapillary OCT circle scans. Statistical comparisons were made using an unpaired Student's t-test. C. Spider plots of NGI, ONL, and total retinal thickness in control (red) and *Sort1-/-* animals (blue) quantified in horizontal line scans through the optic disc. Significance levels: * [0.01, 0.05]; ** [0.001,0.01]; *** [0.0001, 0.001]; **** [0, 0.0001].

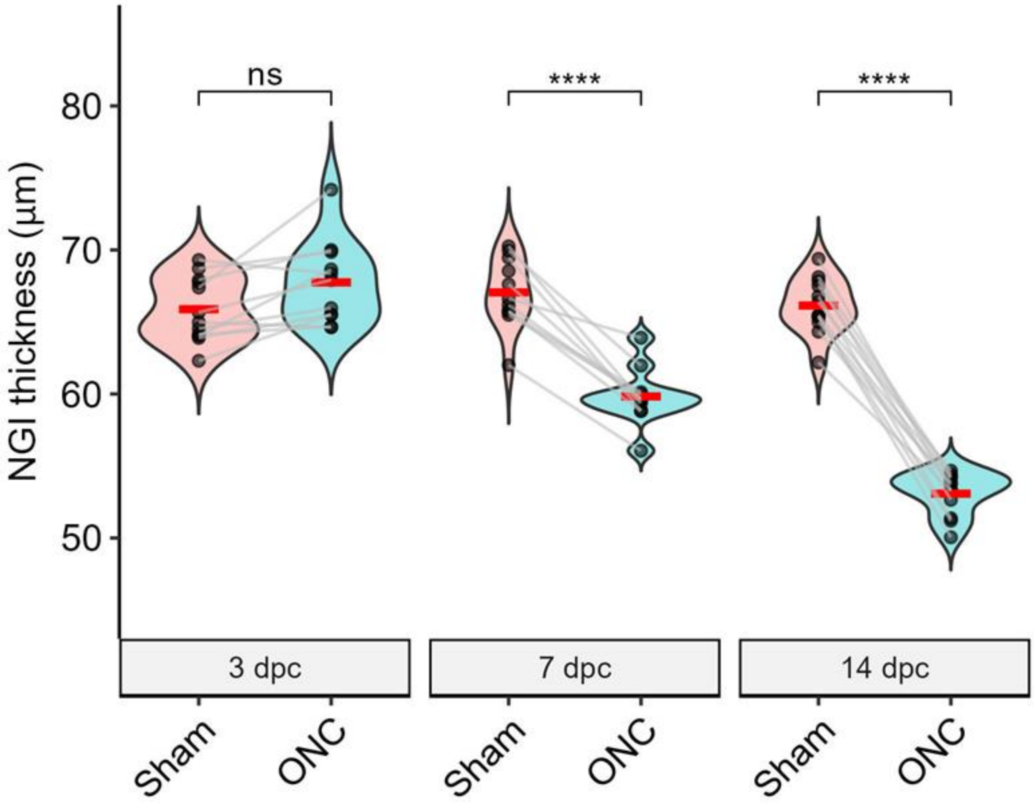

**Figure S4. NGI thickness in animals used for Western blot and RT-qPCR.** Violin plot with quantification of the combined thickness of the retinal nerve fiber, ganglion cell, and inner plexiform layer collectively denoted NGI 3-, 7-, and 14-days post crush in ONC and sham-operated mice. Statistical comparisons were performed using unpaired t-tests. Significance levels: * [0.01, 0.05]; ** [0.001,0.01]; *** [0.0001, 0.001]; **** [0, 0.0001].

**A Sortilin 3 dpc**

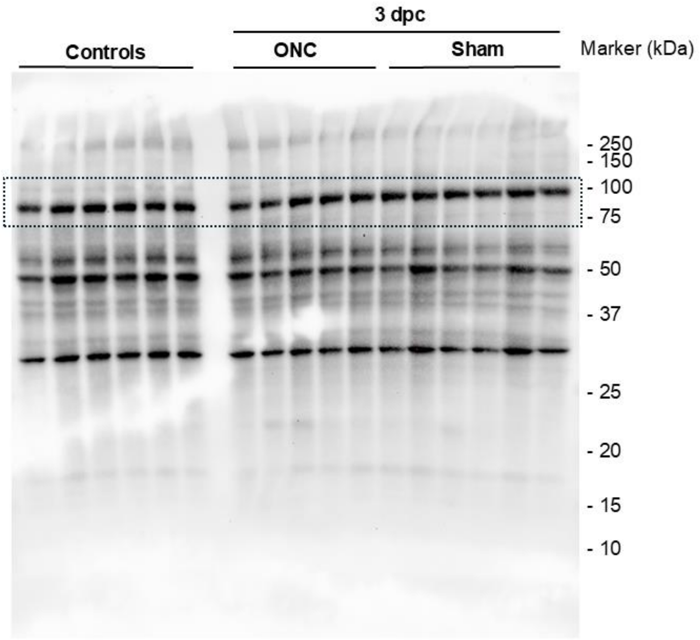

**B Sortilin 7 dpc**

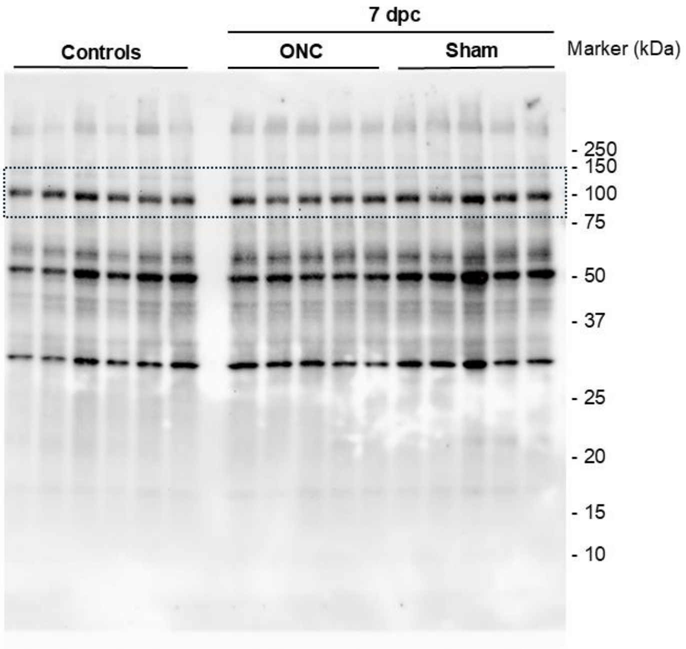

**C Sortilin 14 dpc**

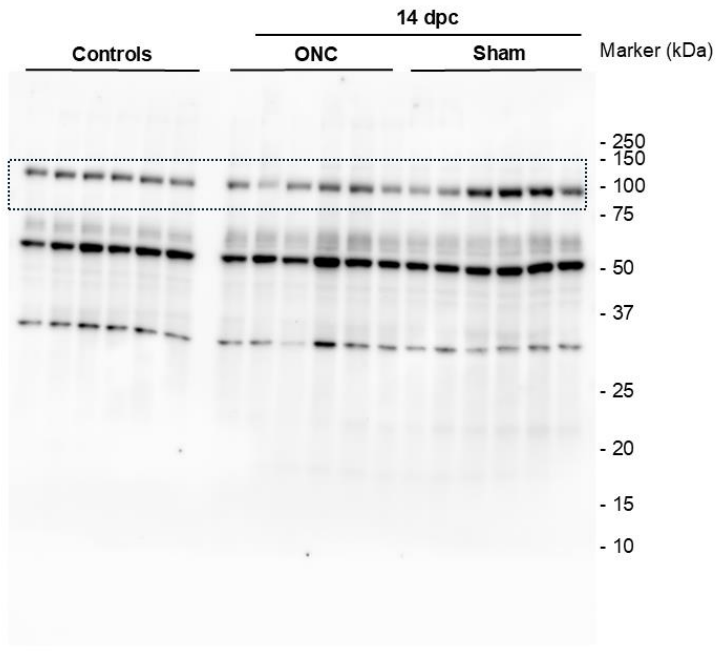

**D p75$^{NTR}$ 3 pdc**

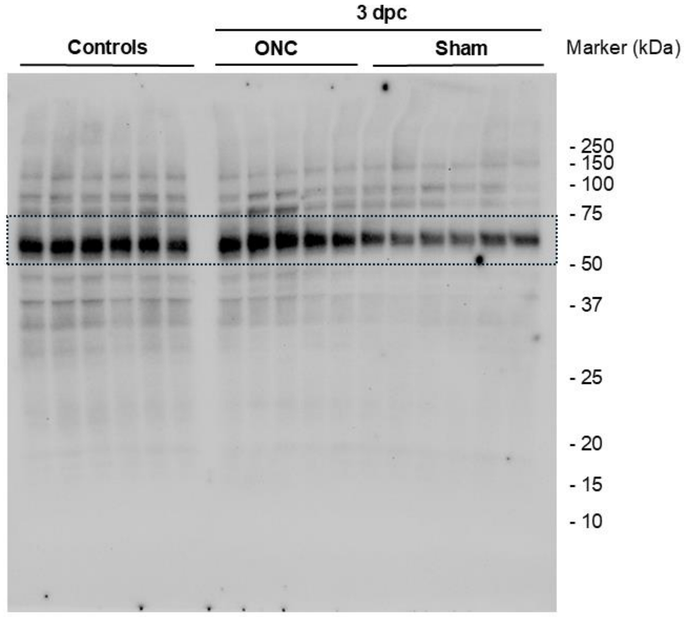

**E p75$^{NTR}$ 7 dpc**

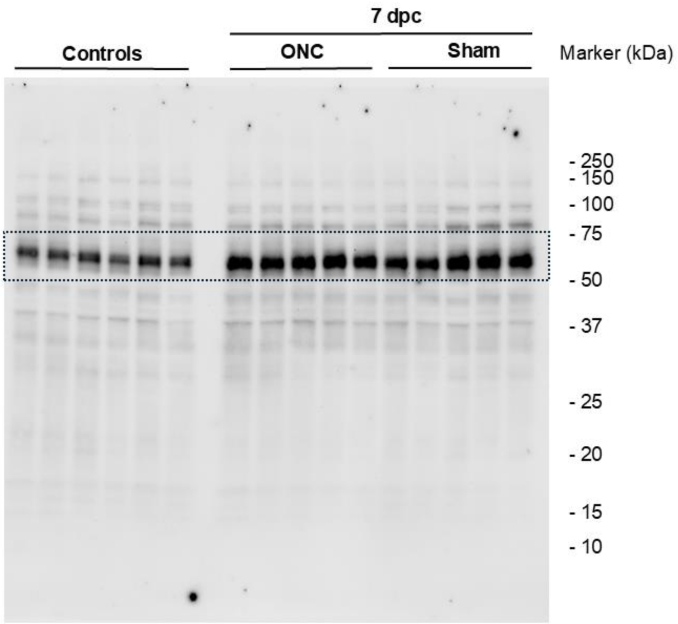

**F p75$^{NTR}$ 14 dpc**

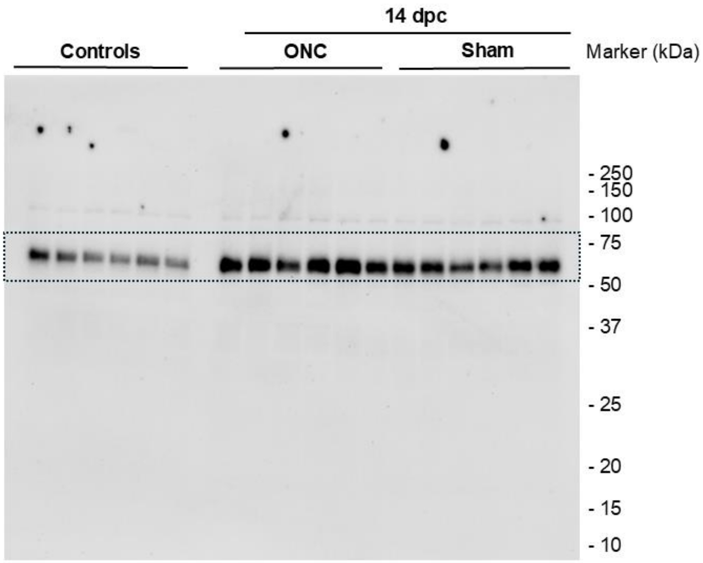

**G Total protein 3 dpc**

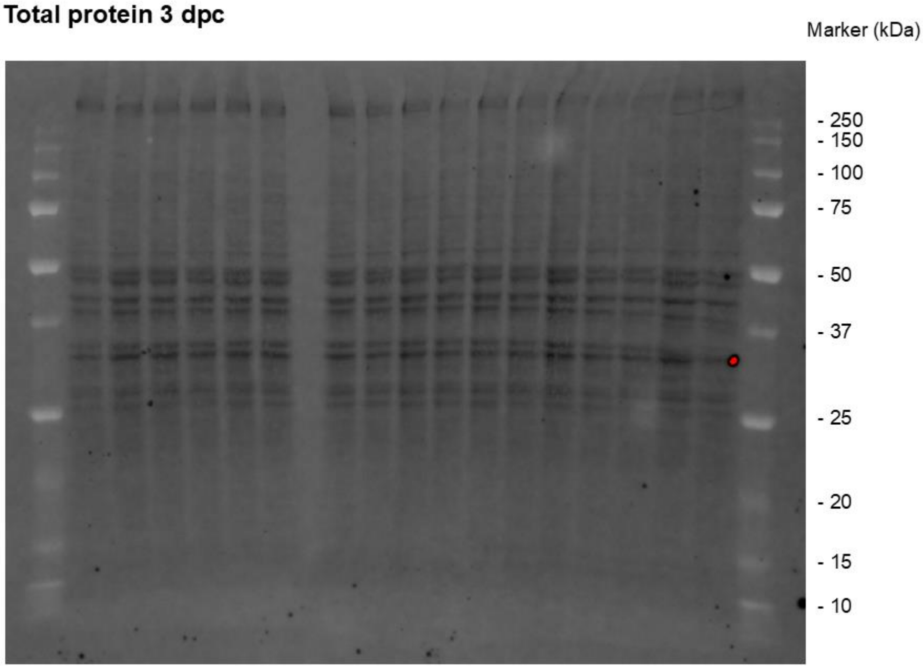

**H Total protein 7 dpc**

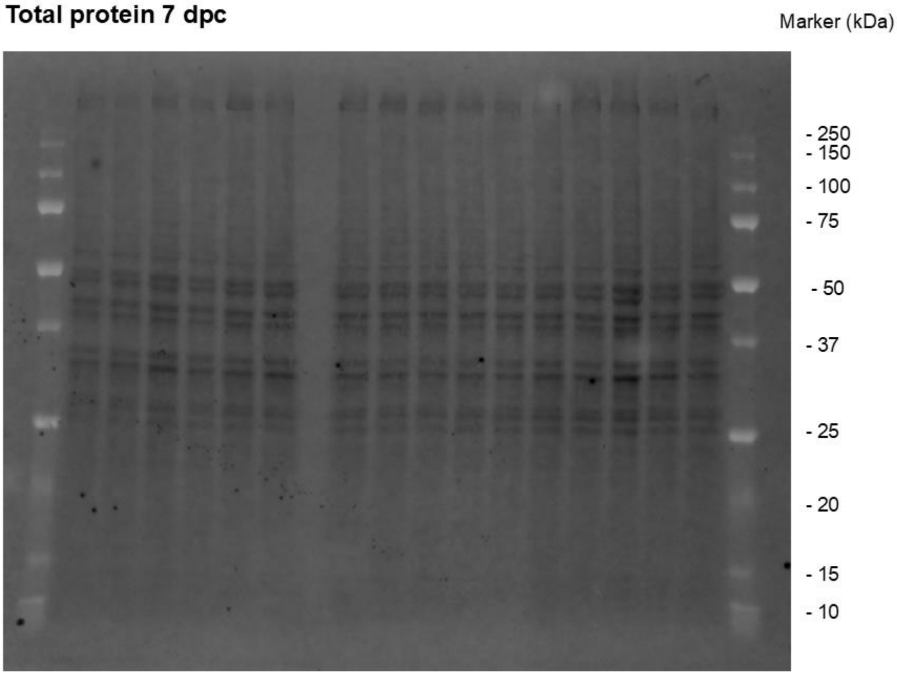

I Total protein 14 dpc

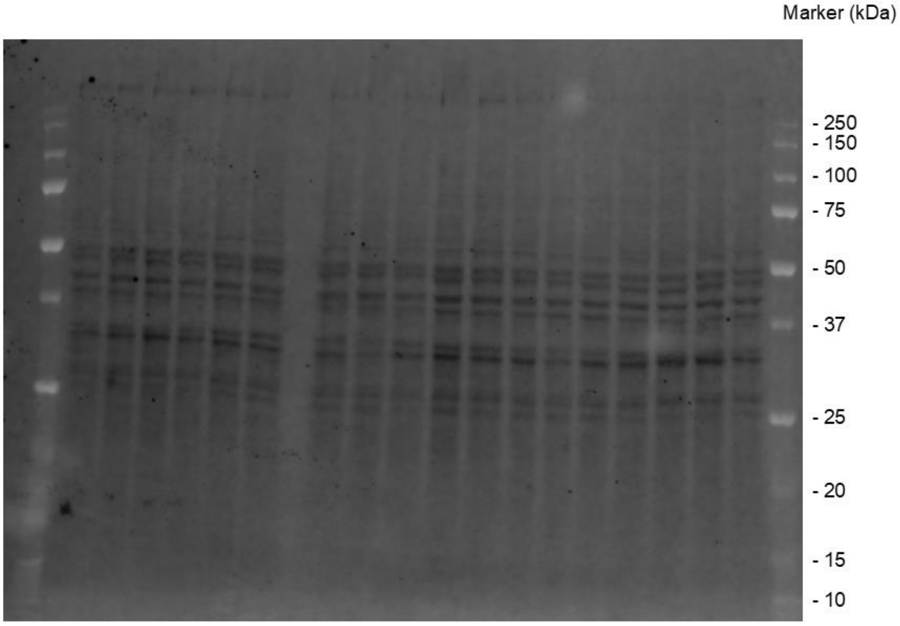

**Figure S5. Western blot analysis for sortilin and p75$^{NTR}$ in retinal tissue lysate following optic nerve crush.** Image shows membrane stained for sortilin (A-C) and p75$^{NTR}$ (D-F) with data cropped for Figure 3E highlighted by dotted boxes. Additional total protein, quantified for use in loading normalization is shown (G-I). dpc, days post crush.

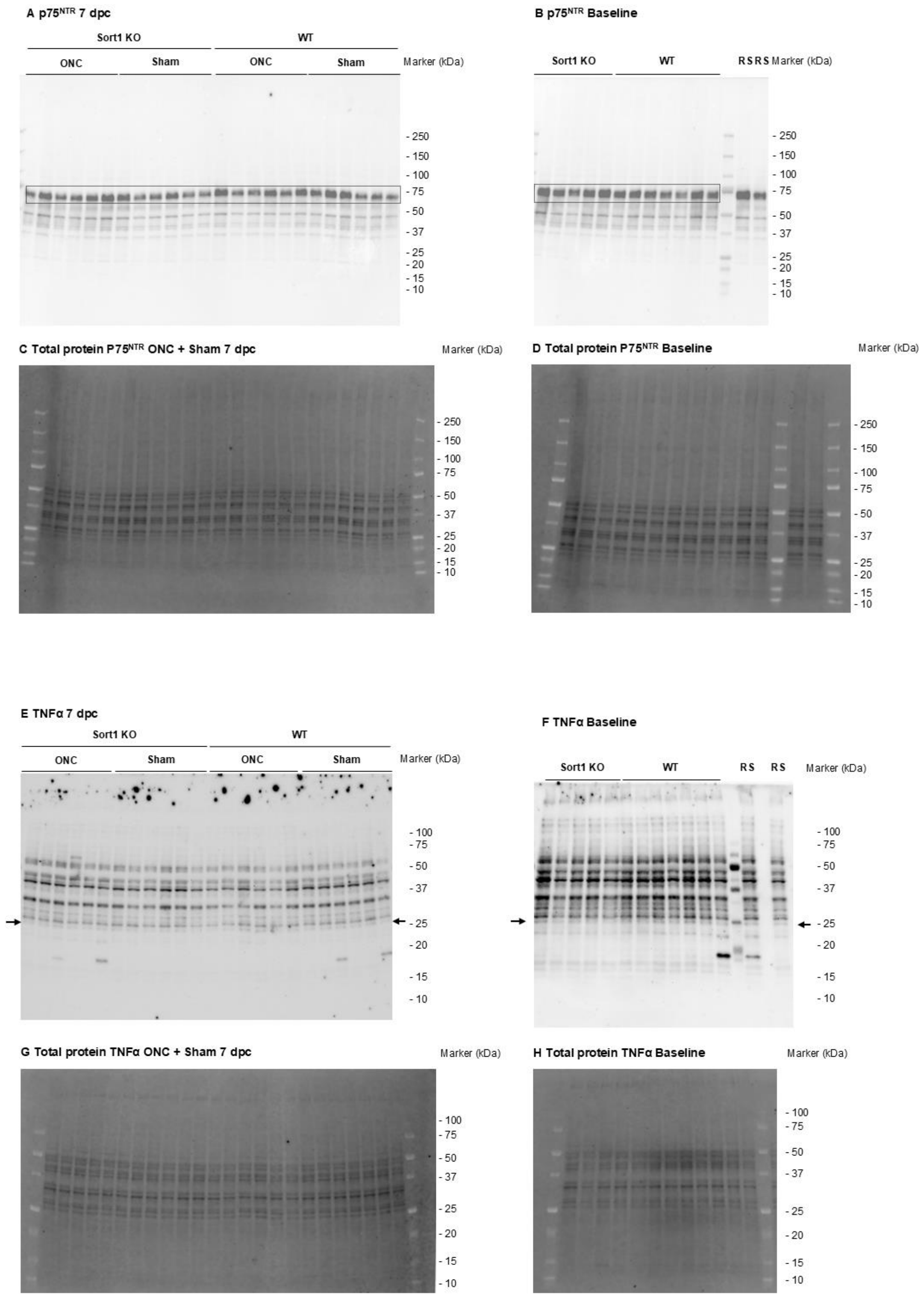

A p75NTR 7 dpc
Sort1 KO
WT
ONC
Sham
ONC
Sham
Marker (kDa)
- 250
- 150
- 100
- 75
- 50
- 37
- 25
- 20
- 15
- 10
B p75NTR Baseline
Sort1 KO
WT
R S R S
Marker (kDa)
- 250
- 150
- 100
- 75
- 50
- 37
- 25
- 20
- 15
- 10
C Total protein P75NTR ONC + Sham 7 dpc
Marker (kDa)
- 250
- 150
- 100
- 75
- 50
- 37
- 25
- 20
- 15
- 10
D Total protein P75NTR Baseline
Marker (kDa)
- 250
- 150
- 100
- 75
- 50
- 37
- 25
- 20
- 15
- 10
E TNFα 7 dpc
Sort1 KO
WT
ONC
Sham
ONC
Sham
Marker (kDa)
- 100
- 75
- 50
- 37
- 25
- 20
- 15
- 10
F TNFα Baseline
Sort1 KO
WT
R S
R S
Marker (kDa)
- 100
- 75
- 50
- 37
- 25
- 20
- 15
- 10
G Total protein TNFα ONC + Sham 7 dpc
Marker (kDa)
- 100
- 75
- 50
- 37
- 25
- 20
- 15
- 10
H Total protein TNFα Baseline
Marker (kDa)
- 100
- 75
- 50
- 37
- 25
- 20
- 15
- 10

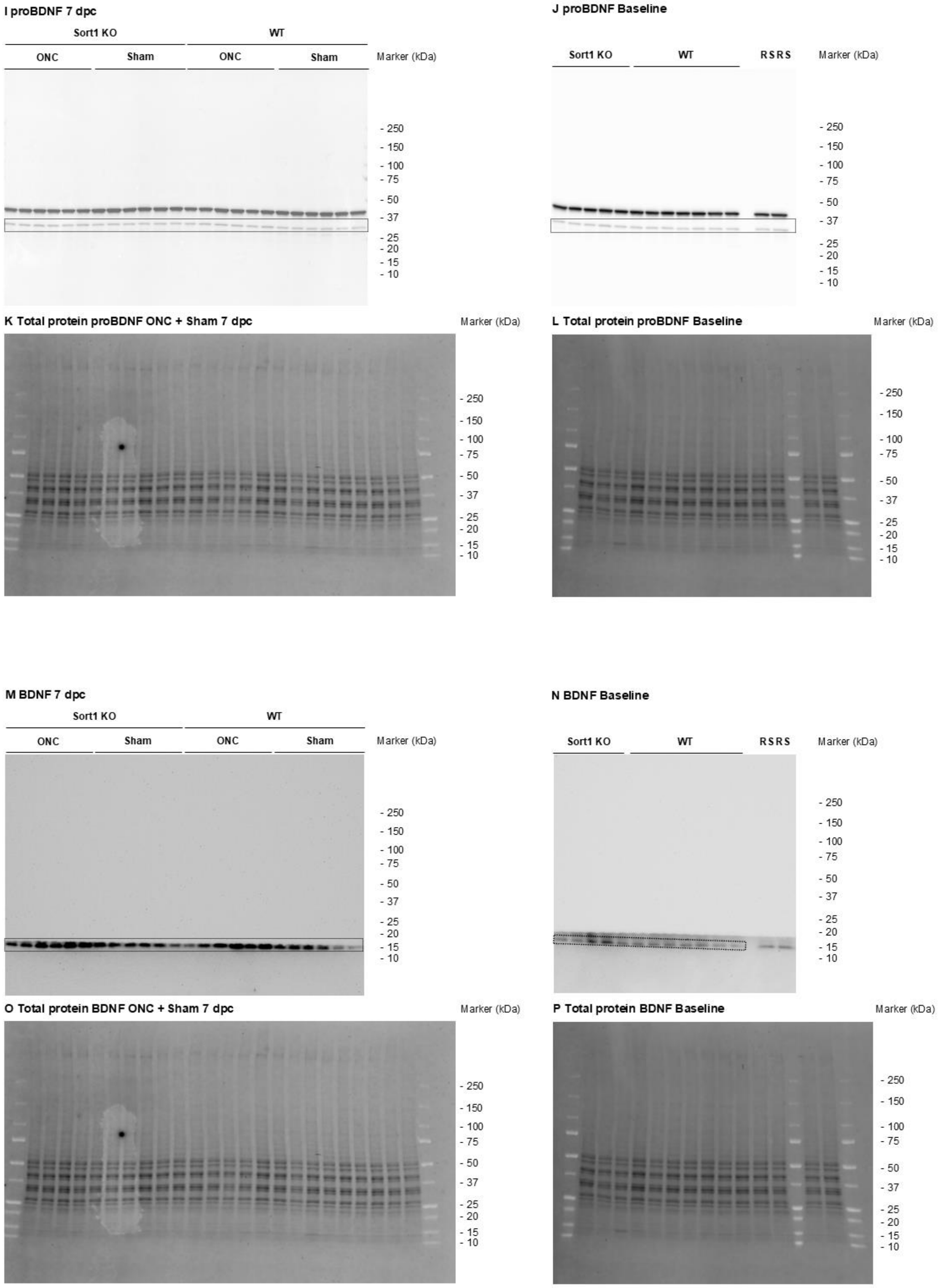
I proBDNF 7 dpc
Sort1 KO
WT
ONC
Sham
ONC
Sham
Marker (kDa)
- 250
- 150
- 100
- 75
- 50
- 37
- 25
- 20
- 15
- 10
J proBDNF Baseline
Sort1 KO
WT
RSRS
Marker (kDa)
- 250
- 150
- 100
- 75
- 50
- 37
- 25
- 20
- 15
- 10
K Total protein proBDNF ONC + Sham 7 dpc
Marker (kDa)
- 250
- 150
- 100
- 75
- 50
- 37
- 25
- 20
- 15
- 10
L Total protein proBDNF Baseline
Marker (kDa)
- 250
- 150
- 100
- 75
- 50
- 37
- 25
- 20
- 15
- 10
M BDNF 7 dpc
Sort1 KO
WT
ONC
Sham
ONC
Sham
Marker (kDa)
- 250
- 150
- 100
- 75
- 50
- 37
- 25
- 20
- 15
- 10
N BDNF Baseline
Sort1 KO
WT
RSRS
Marker (kDa)
- 250
- 150
- 100
- 75
- 50
- 37
- 25
- 20
- 15
- 10
O Total protein BDNF ONC + Sham 7 dpc
Marker (kDa)
- 250
- 150
- 100
- 75
- 50
- 37
- 25
- 20
- 15
- 10
P Total protein BDNF Baseline
Marker (kDa)
- 250
- 150
- 100
- 75
- 50
- 37
- 25
- 20
- 15
- 10

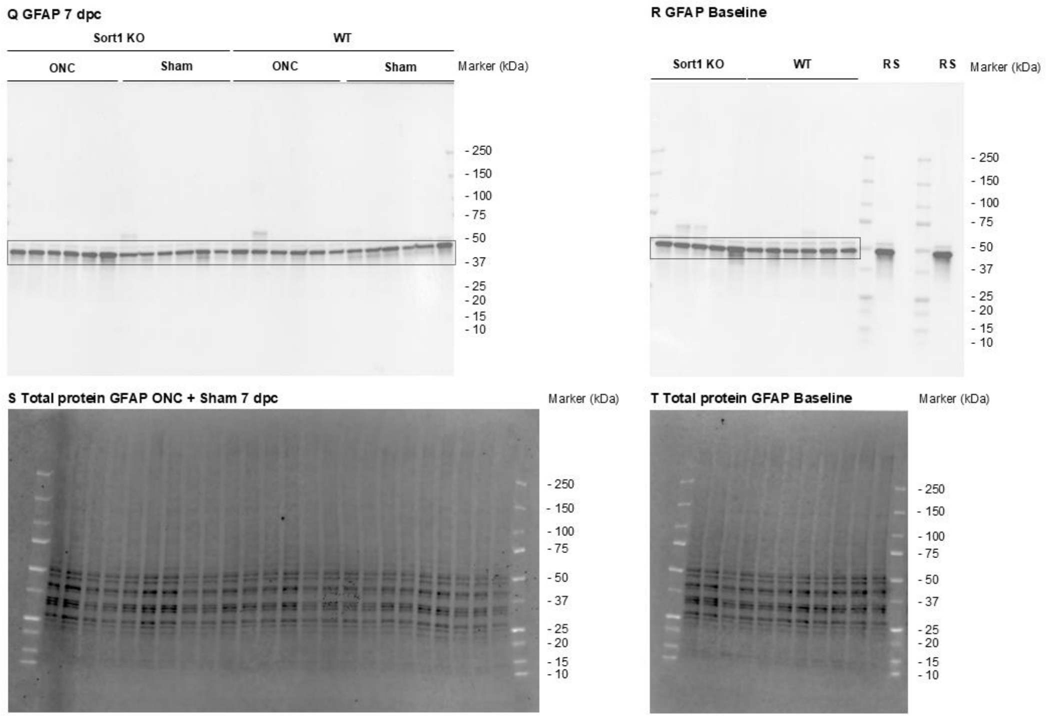

**Figure S6. Western blot analysis for p75$^{NTR}$, TNFα, BDNF, and GFAP in retinal tissue lysate following optic nerve crush.** Image shows membrane stained for p75$^{NTR}$ (A-D), TNFα (E-H), proBDNF (I-L), BDNF (M-P) and GFAP (Q-T) with data cropped for Figure 4E highlighted by dotted boxes or arrows. Baseline levels of each of the analyzed proteins are shown in B, F, J, N, and R. To enable specific detection of the BDNF signal (M,N), portions of the membrane were masked to prevent interference from nonspecific binding. Additional total protein, quantified for use in loading normalization is shown (C, D, G, H, K, L, O, P, S, and T). dpc, days post crush; RS, reference sample for use in inter-gel normalization.

#### Supplementary Table 1: RT-qPCR primers

| | |
|---|---|
| *Sort1* forward | 5’-CACATGGAGCATGGCACAAC-3’ |
| *Sort1* reverse | 5’-ACCCGGTATCTCCAGGTTCA-3’ |
| *Ngfr* forward | 5’- CCCTGCCTGGACAGTGTTAC -3’ |
| *Ngfr* reverse | 5’- ACAGGGAGCGGACATACTCT -3’ |
| *Gapdh* forward | 5’-GGTGAAGGTCGGTGTGAACG-3’ |
| *Gapdh* reverse | 5’-CTCGCTCCTGGAAGATGGTG-3’ |

Primers were purchased from Eurofins Genomics (Ebersberg, Germany)
